# Druggable genome screen identifies new regulators of the abundance and toxicity of ATXN3, the Spinocerebellar Ataxia Type 3 disease protein

**DOI:** 10.1101/690818

**Authors:** Naila S. Ashraf, Joanna R. Sutton, Yemen Yang, Bedri Ranxhi, Kozeta Libohova, Emily D. Shaw, Anna J. Barget, Sokol V. Todi, Henry L. Paulson, Maria do Carmo Costa

**Affiliations:** Department of Neurology, Michigan Medicine, University of Michigan, Ann Arbor, MI, USA; Department of Pharmacology, Wayne State University School of Medicine, Detroit, MI, USA; Department of Neurology, Wayne State University School of Medicine, Detroit, MI, USA

**Keywords:** polyglutamine, spinocerebellar ataxia, Machado-Joseph disease, neurodegeneration, high-throughput screen, human embryonic stem cells, *Drosophila*

## Abstract

**Background:** Spinocerebellar Ataxia type 3 (SCA3, also known as Machado-Joseph disease) is a neurodegenerative disorder caused by a CAG repeat expansion encoding an abnormally long polyglutamine (polyQ) tract in the disease protein, ataxin-3 (ATXN3). No preventive treatment is yet available for SCA3. Because SCA3 is likely caused by a toxic gain of ATXN3 function, a rational therapeutic strategy is to reduce mutant ATXN3 levels by targeting pathways that control its production or stability. Here, we sought to identify genes that modulate ATXN3 levels as potential therapeutic targets in this fatal disorder.

**Methods:** We screened a collection of siRNAs targeting 2742 druggable human genes using a cell-based assay based on luminescence readout of polyQ-expanded ATXN3. From 317 candidate genes identified in the primary screen, 100 genes were selected for validation. Among the 33 genes confirmed in secondary assays, 15 were validated in an independent cell model as modulators of pathogenic ATXN3 protein levels. Ten of these genes were then assessed in a *Drosophila* model of SCA3, and one was confirmed as a key modulator of physiological ATXN3 abundance in SCA3 neuronal progenitor cells.

**Results:** Among the 15 genes shown to modulate ATXN3 in mammalian cells, orthologs of *CHD4*, *FBXL3*, *HR* and *MC3R* regulate mutant ATXN3-mediated toxicity in fly eyes. Further mechanistic studies of one of these genes, *FBXL3*, encoding a F-box protein that is a component of the SKP1-Cullin-F-box (SCF) ubiquitin ligase complex, showed that it reduces levels of normal and pathogenic ATXN3 in SCA3 neuronal progenitor cells, primarily via a SCF complex-dependent manner. Bioinformatic analysis of the 15 genes revealed a potential molecular network with connections to tumor necrosis factor-*α*/nuclear factor-kappa B (TNF/NF-kB) and extracellular signal-regulated kinases 1 and 2 (ERK1/2) pathways.

**Conclusions:** We identified 15 druggable genes with diverse functions to be suppressors or enhancers of pathogenic ATXN3 abundance. Among identified pathways highlighted by this screen, the FBXL3/SCF axis represents a novel molecular pathway that regulates physiological levels of ATXN3 protein.

## Introduction

The polyglutamine (polyQ) diseases are inherited neurodegenerative diseases caused by expanded CAG repeats that encode abnormally long glutamine repeats in the disease proteins [1, 2]. Spinocerebellar Ataxia type 3 (SCA3) is one of nine known polyQ disorders and the most common dominant ataxia, primarily manifesting with degeneration of the cerebellum, brainstem, spinal cord, and basal ganglia [3–7]. The CAG repeat in the *ATXN3* gene, which normally is 12 to 44 triplets, becomes expanded to ~60 to 87 repeats in SCA3 [8, 9]. Despite sharing a propensity to misfold and aggregate, polyQ disease proteins differ in size, cellular localization and biological function. Moreover, polyQ disorders show distinctive symptomatology and neuropathology, indicating that the specific protein context in which expanded polyQ is embedded influences the pathogenic mechanisms in each disease [2].

While many advances have been made in understanding pathomechanisms and promising therapies may be on the horizon for polyQ diseases, no disease-modifying treatments exist yet. Reducing levels of mutant *ATXN3* transcripts and/or protein using nucleotide-based approaches or small molecules has been reported by us and others as an encouraging therapeutic strategy for SCA3 [10–19]. Another route to suppressing polyQ disease protein abundance in the mammalian brain is to manipulate specific pathways used by cells to control mutant protein production, stability, or clearance. Unfortunately, the mechanisms underlying cellular handling of ATXN3 and other mutant polyQ proteins remain poorly understood.

Here, we carried out an unbiased druggable genome siRNA screen in a cell-based assay to identify genes and pathways that modulate levels of expanded-polyQ ATXN3. Downstream validation of identified genes was then performed in *Drosophila* models of SCA3 and neuronal progenitor cells (NPCs) derived from human embryonic stem cells (hESCs) harboring an expanded CAG repeat in *ATXN3*. We identified novel genes that regulate ATXN3 levels in mammalian cells and modulate mutant ATXN3-mediated toxicity in *Drosophila*, suggesting new therapeutic targets in SCA3 and perhaps similar disorders.

## Materials and Methods

### Druggable genome siRNA primary screen

High-throughput screens were carried out at the University of Michigan Center for Chemical Genomics (CCG). We screened the druggable genome subset of the human siGENOME siRNA SMARTpool library (Dharmacon) targeting 2742 genes. Each library stock plate comprising 280 siRNA SMART-pools targeting druggable genes and four internal library siRNA controls for viability at 500 nM was screened in triplicate by reverse transfection of stably transfected 293/ATXN3Q81:FF-Luc (ATXN3-Luc) cells. Sample siRNAs were screened at a final concentration of 50 nM and control siGENOME siRNAs (RISC-Free Non-targeting, and SMARTpools for ATXN3 and BECLN1) at 20 nM. siRNAs (4 μL) were transferred from each library stock plate (500 nM) and a control stock plate (200 nM) to three 384-well black tissue culture-treated assay plates (Greiner-Bio CellStar) using Biomek FX (Beckman Coulter, Fullerton, CA) (Supplementary Figure 1). Lipofectamine RNAiMAX transfection reagent (Thermo Fisher Scientific) was diluted in OPTIMEM (Thermo Fisher Scientific) (0.1 μL of RNAiMax + 5.9 μL of OPTIMEM) and 6 μL of the mixture were added per well to the assay plates containing the siRNAs using an automated dispenser (Multidrop Combi, Thermo Fisher). Plates were briefly centrifuged and incubated at room temperature (RT) for 30 min. ATXN3-Luc cells in DMEM/ 10% fetal bovine serum were added to the assay plates (10^4^ cells in 30 μL per well) and briefly centrifuged. Cell lysis buffer (Promega) (10 μL) was added to wells A1, A24, P1 and P24 (positive controls for viability) and plates were incubated at 37°C, 5% CO_2_ for 48 hours. After incubation, medium was aspirated leaving about 10 μL per well, and 5 μL of CellTiter-Fluor Cell Viability Assay reagent (Promega) was added to each well. Plates were incubated at 37°C, 5% CO_2_ for 1 hour and viability was assessed by fluorescence reading (Exc 390/ Em 505 nm) at the PHERAstar (BMG Labtech). Ten μL of Steady-Glo Luciferase Assay System were subsequently added to each well, and after 10 min incubation at room temperature, the activity of firefly Luciferase was measured by PHERAstar (BMG Labtech). Data was uploaded and computed by MScreen Database [20] for negative and positive controls on a plate by plate basis. Percent viability for sample siRNAs was calculated based on Lysed cells (0%) and siRNA control RISC-Free Nontargeting (100%). The resulting percent luminescence activity for the siRNAs was computed relative to the negative siRNA control (RISC-Free Nontargeting) at 0% and the positive siRNA control (targeting ATXN3) at 100%.

### siRNA confirmation screens

A customized library of four individual siGENOME siRNAs per gene for 100 genes was screened in parallel in ATXN3-Luc and Luc (counter-screen) cells in triplicate using the above protocol. Genes for which at least one individual siRNA showed selective activity on the ATXN3-Luc assay and not on the Luc-assay were considered for follow-up validation.

### siRNA reverse transfection of stably transfected HEK293 cells

HEK293 cells stably expressing FLAG-ATXN3Q80 [21] were cultured in DMEM/ 10% FBS/ 1% Penicillin-Streptomycin. Sample individual siGENOME siRNAs were screened at 50 nM and control siRNAs (RISC-Free Non-targeting, and SMARTpool for ATXN3) at 20 nM in 24-well plate setups. Briefly, 40 μL of siRNA were incubated with 60 μL of transfection mixture (1 μL of RNAiMax + 59 μL of OPTIMEM) for 30 min at room temperature, after which 10^5^ cells in 300 μL of medium were added per well. Cells were incubated at 37°C, 5% CO_2_ for 48 hours.

### Generation and culture of human control and SCA3 neuronal progenitor cells (NPCs)

Control and SCA3 human embryonic stem cells (hESCs), respectively, UM4-6 NIH registry #0147 and UM134-1 PGD NIH registry #0286), were acquired from the MStem Cell Lab, University of Michigan. All experiments using hESCs were previously approved by the Human Pluripotent Stem Cell Research Oversight (HPSCRO) of the University of Michigan (Application 1097). Control and SCA3 NPCs were generated from the respective hESC lines (passages 18 (P18) and 15 (P15), respectively) using the STEMdiff Neural System (STEMCELL Technologies). Briefly, 2 ×10^6^ SCA3 hESCs were resuspended in 2 mL of STEMdiff Neural Induction Medium (NIM)/ SMADi/ 10 μM Y-27632 and plated in one well of a 6-well plate coated with poly-L-ornithine (PLO) and laminin (lam) and incubated at 37°C, 5% CO_2_. Daily full medium changes were performed with NIM/ SMADi until Day 6, when cells (80-90% confluent) were passaged into two 60 mm plates coated with PLO/lam (P1) using ACCUTASE and NIM/ SMADi/ 10 μM Y-27632. Medium was changed daily and cells were passaged as above at Day 10 (P2). At Day 13 (P3), cells were passaged and plated in STEMdiff Neural Progenitor Medium (NPM) and since then cells were expanded and maintained in NPM. Control and SCA3 NPCs (P4) were evaluated for expression of NPC markers and cells of passage four or higher were frozen in STEMdiff Neural Progenitor Freezing Medium in liquid nitrogen. Cells were thawed and re-expanded as needed for experiments.

### Electroporation of SCA3 neuronal progenitor cells (NPCs)

Approximately 10^6^ SCA3 NPCs (P8-P11) were electroporated with 1 µg of pCMV-SPORT6.FBXL3 (Dharmacon, MHS6278-202759846) or 1 µg of pCMV-SPORT6.FBXL3 and 40 nM ON-TARGETplus human CUL1 (8454) siRNA SMARTpool (Dharmacon, L-004086) using the Neon transfection system (Invitrogen) following the manufacturer instructions and 1400 V, 20 ms, 2 pulses per condition. Cells were plated in 1.5 mL of NPM of a 12-well plate and incubated at 37°C, 5% CO_2_. Medium was changed after 32 hours either with NPM or NPM/ 2 µM MLN4924 and cells were incubated for 16 additional hours. Forty-eight hours after transfection, cells were collected in 200 µL of RIPA buffer/ COMPLETE (Roche Diagnostics)/ PhosSTOP (Sigma) for protein extraction or 200 µL of RLT buffer (Qiagen) for RNA extraction and stored at −80°C until further sample processing.

### Western Blotting

Total proteins from mammalian cells were extracted from cells by resuspension and homogenization in RIPA buffer containing protein inhibitors (Complete, Roche Diagnostics) and phosphatase inhibitors (Phospho-STOP, Sigma), followed by sonication and centrifugation at 4°C. Supernatants (soluble proteins) were collected and total proteins were quantified using the BCA method (Pierce). For fly-based samples, 10 dissected heads per group were mechanically homogenized in boiling 2% SDS lysis buffer, sonicated, boiled for 10 minutes and centrifuged at 14,400 X g before loading. Total protein lysates (20 μg) were resolved in 10% SDS-PAGE gels, and corresponding PVDF membranes were incubated overnight at 4°C with primary antibodies: mouse anti-ATXN3 (1H9) (1:2000, MAB5360, Millipore), rabbit anti-MJD [22] (1:20000), rabbit anti-FBXL3 (1:500, ab96645, Abcam), rabbit anti-CUL1 (EPR3103Y) (1:1000, ab75817, Abcam), and rabbit anti-α-Tubulin (11H10) (1:10000, #2125, Cell Signaling Technology). Primary antibodies were detected by incubation with peroxidase-conjugated anti-mouse and anti-rabbit antibodies (1:10000, Jackson Immuno Research Laboratories) followed by reaction with ECL-Plus reagent (Western Lighting, PerkinElmer) and exposure to autoradiography films. Film band intensity was quantified by densitometry by Image J.

### Immunofluorescence

PLO/lam-coated coverslips with SCA3 NPCs were washed with PBS, fixed with 4% paraformaldehyde/ PBS for 15 min, washed three times with PBS and stored at 4°C until further processing. Cells were permeabilized with 0.5% Triton X-100/ PBS for 20 min, washed with Tween-20/ PBS (PBS-T), blocked in 5% goat serum/ PBS for one hour, and incubated overnight at 4°C with primary antibodies diluted in 5% goat serum/ PBS: rabbit anti-PAX6 (1:250, 60433S, Cell Signaling), rabbit anti-SOX1 (1:1000, 4194S, Cell Signaling), mouse anti-NESTIN (1:250, 33475S, Cell Signaling), rabbit anti-MJD (1:1000), and mouse anti-ATXN3 (1H9) (1:250, MAB5360, Millipore). Cells were washed with PBS-T, incubated with corresponding secondary antibodies goat anti-rabbit and anti-mouse conjugated with Alexa Fluor 488 or 568 (1:1000, Invitrogen) diluted in 5% goat serum/ PBS for 1h, incubated with DAPI for 10 min, and washed with PBS-T. Immunostained coverslips were then mounted in slides using Prolong Gold medium (Invitrogen) and saved at 4°C until imaged on a Nikon A1 high sensitivity confocal microscope.

### RNA extraction and quantitative RT-PCR

Total RNA from cells was extracted using the RNeasy mini kit (Qiagen) following manufacturer’s instructions. Reverse transcription of 1 μg of total RNA per sample was performed using the iScript cDNA synthesis kit (Bio-RAD). Transcript levels were determined by quantitative real-time RT-PCR as previously reported [14] using primers provided in Supplementary Table 1 and normalizing expression to *ATCB* transcript levels.

### *Drosophila* experiments

For all stocks and for experimental procedures, adult males and virgin females were crossed, raised and maintained at 25°C under diurnal conditions in standard cornmeal media. All examined flies were heterozygous for driver and transgenes. Once offspring emerged from pupal cases, they were aged under the same conditions described above for seven days, at which time heads were dissected and imaged using an Olympus BX53 microscope equipped with a DP72 digital camera for GFP fluorescence experiments, or whole flies were fixed and processed for histological sections, described below. Fluorescence from each eye was quantified using the publicly available ImageJ software. Average retinal fluorescence for each treatment condition was calculated as previously described [23–27]. RNAi fly lines used for the work described in this manuscript are listed in Supplementary Table 2. The GMR-Gal4 driver (#8605) and UAS-mCD8-GFP (#5137) were from the Bloomington *Drosophila* Stock center. The UAS-ataxin-3Q77 line has been described before [13, 28]. For histological sections, adult fly wings and proboscises were removed, and flies were fixed in 2% glutaraldehyde/ 2% paraformaldehyde in Tris-buffered saline with 0.1% Triton X-100. The fixed flies were dehydrated in a series of 30%, 50%, 75%, and 100% ethanol, and 100% propylene oxide. Dehydrated specimens were embedded in Poly/Bed812 (Polysciences) and fly heads were sectioned at 5 µm. Sectioned heads were stained with toluidine blue.

### Bioinformatic analysis

Gene lists were analyzed for biological functions and network analysis was performed with the Ingenuity Pathway Analysis software (Qiagen) using the whole-human genome as the reference gene set and known direct and indirect gene relationships.

### Statistical analysis

Levels of proteins and transcripts, and fluorescence in *Drosophila* were compared using one-tailed or two-tailed student’s t-test. A *P*<0.05 was considered statistically significant for all analyses. Data were analyzed using IBM SPSS Statistics 22.

## Results

### Mammalian cell-based siRNA screen identifies modulators of pathogenic ATXN3

To identify genes that regulate levels of pathogenic ATXN3 in mammalian cells, we used our previously developed ATXN3-Luc cellular assay (Figure 1A,B) [13] to screen the druggable genome subset of the human siGENOME siRNA library (Dharmacon). This library comprises SMARTpools of four individual siRNAs targeting 2742 genes that are considered potential therapeutic targets, including G-protein coupled receptors (GPCRs), ion channels, protein kinases, proteases, phosphatases and ubiquitin conjugation enzymes (https://dharmacon.horizondiscovery.com/rnai/). The ATXN3-Luc assay measures chemiluminescence in HEK293 cells stably overexpressing FLAG-tagged human ATXN3 harboring an expanded polyQ repeat in the disease range (Q81) fused to firefly Luciferase, under the control of a CMV promoter (Figure 1A,B) [13]. We reasoned that by screening for steady state levels of the ATXN3/Luciferase fusion protein (reported as chemiluminescence), and because its expression is driven by the CMV promoter, we would identify genes that regulate ATXN3 abundance at post-transcriptional steps (e.g. mRNA stability/degradation, and protein translation, folding and turnover). To circumvent false-positive siRNAs that interfere with CMV promoter activity or Luciferase itself, we developed a counter-screen assay for use in confirmation/ validation screens: HEK293 cells stably overexpressing firefly Luciferase controlled by the CMV promoter (Luc assay) (Figure 1A,B).

**Figure 1.**
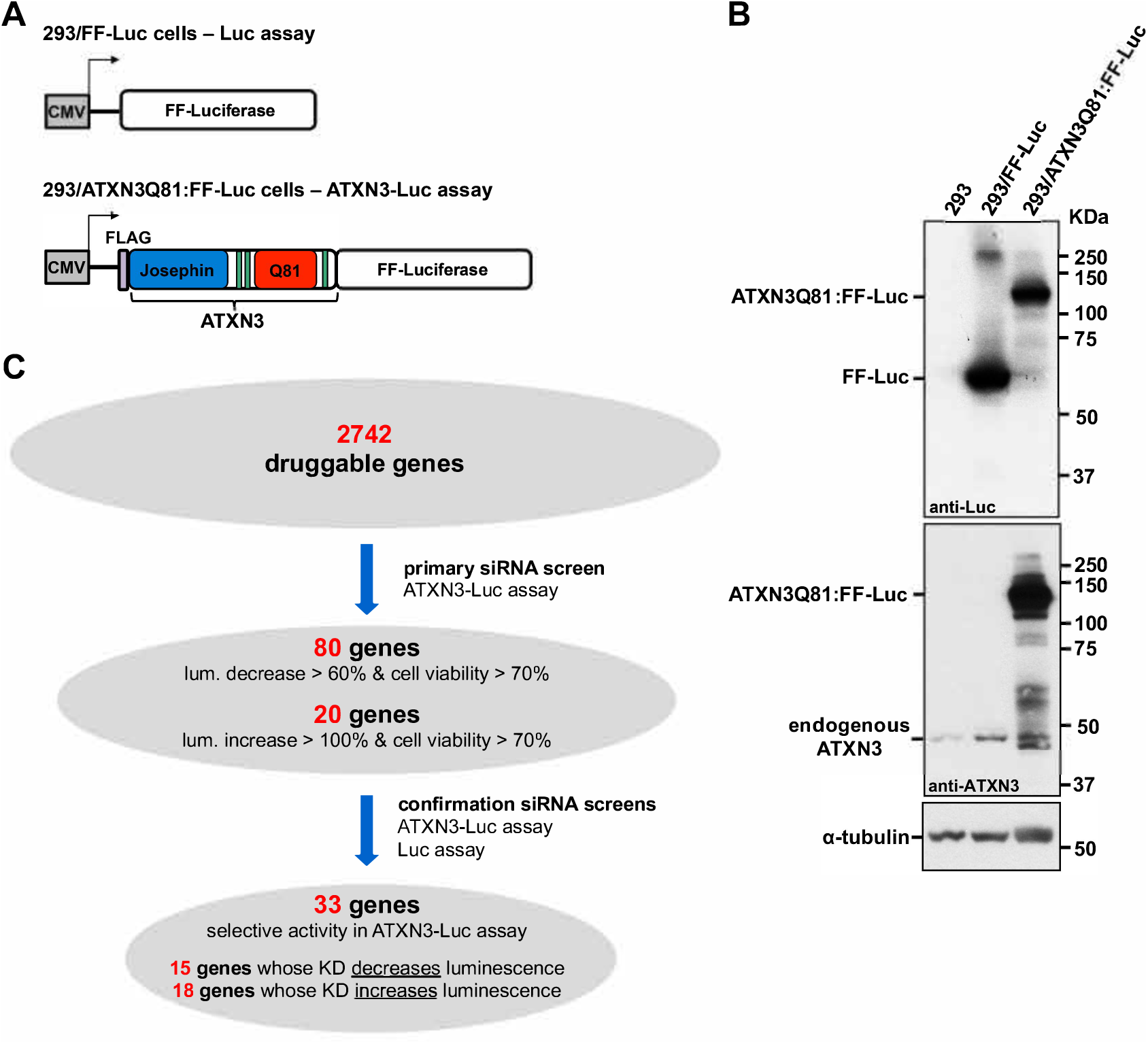
Unbiased cell-based siRNA screen of the druggable genome identifies novel modulators of ATXN3 levels. A) Schematic of constructs expressing firefly Luciferase by itself (Luc assay) or FLAG-tagged human ATXN3Q81 fused to Luciferase (ATXN3-Luc assay) in stably expressing HEK293 cell lines. FF: firefly. B) Western blots with anti-ATXN3 (1H9) and anti-Luciferase (Luc) antibodies show expression of ATXN3Q81:FF-Luc fusion protein and FF-Luc, respectively, in ATXN3-Luc and Luc assays. C) Summary of the iterative screens using the ATXN3-Luc and Luc assays to select 33 genes that modulate levels of mutant ATXN3 for subsequent studies.

We performed the primary screen in ATXN3-Luc cells, employing a 384-well plate format with pools of four individual siRNAs targeting a single gene per well. Pooled siRNAs were tested in triplicate plates for their efficacy to decrease or increase levels of luminescence and to affect cell viability (Figure 1C). All plates included built-in controls for luminescence and cell viability readouts (Supplementary Figure 1): 1) siGENOME RISC-Free Non-Targeting was used as a negative control for both readouts; 2) pooled siGENOME siRNAs targeting ATXN3 were used as a positive control for suppressors of luminescence; 3) because BECN1 was shown to clear ATXN3 in SCA3 mouse models [29, 30], pooled siGENOME siRNAs against BECN1 were used as positive control for enhancers of luminescence; and 4) cell lysis buffer was used as a positive control for suppressors of cell viability.

A total of 33 plates were screened in two assays, showing an average plate Z factor of 0.82 for luminescence assessment. Some hits that reduce luminescence may be false positives due to cell death caused by depletion of an essential gene; thus, to identify genes whose knockdown decreases levels of ATXN3 with minimal cell toxicity, we only considered hits that showed cell viability higher than 70% relative to controls. siRNA pools for 317 genes passed this viability cutoff and led to statistically significant increased (N=163) or decreased (N=154) luminescence of at least 50% relative to negative control.

These 317 identified candidate genes were similarly distributed throughout the screened protein families: kinases (N=73 of 675, 10.8%), followed by peptidases (N=44 of 412, 10.7%), G-protein coupled receptors (N=43 of 364, 11.8%), proteins with other functions (N=41 of 271, 15.1%), ion channels (N=38 of 304, 12.5%), enzymes (N=32 of 311, 10.3%), phosphatases (N=30 of 247, 12.1%), transcription regulators (N=10 of 89, 11.9%), transmembrane receptors (N=3 of 39, 7.7%), transporters (N=2 of 29, 7.7%), and growth factors (N=1 of 2, 50%) (Supplementary Figure 2). Analysis of subcellular localization of these 317 genes showed that they are mainly distributed through the cytoplasm (N=116), plasma membrane (N=111) and nucleus (N=51) (Supplementary Figure 2).

Among these 317 hits, we selected 100 genes for confirmation: the top 80 genes whose knockdown decreased luminescence by at least 60%, and the top 20 genes whose knockdown increased luminescence by at least 100% (Figure 1C). We chose to select more genes whose knockdown reduced luminescence because our primary goal is to identify therapeutically compelling targets, and it is more feasible to knock down or suppress the activity of a modifier gene as a therapeutic approach. In the confirmation screens, we assessed in parallel the four individual siGENOME siRNAs per gene in ATXN3-Luc and Luc cells (Figure 1C). For 33 of the 100 genes, we confirmed that at least one siRNA selectively and significantly modulated luminescence levels in ATXN3-Luc cells (Supplementary Table 2): 15 were genes whose knockdown decreased luminescence, and 18 were genes whose knockdown increased luminescence (Figure 1C).

### Fifteen genes confirmed to regulate ATXN3 levels in an independent SCA3 cell line

We next tested whether knockdown of these 33 genes (Supplementary Table 2) modulated levels of ATXN3 in an independent HEK293 cell model stably overexpressing FLAG-tagged human ATXN3 with a polyQ repeat of 80 (ATXN3Q80 cells) [21]. Cells were transiently transfected individually with each of the four siRNAs targeting an identified gene. The efficiency of transcript depletion was confirmed by quantitative RT-PCR (Supplementary Figures 3 and 4), and levels of ATXN3 protein were assessed by Western blot (Figures 2 and 3). In this secondary screen, we considered a gene validated as a modifier of ATXN3 abundance if at least two of the four siRNAs altered levels of expanded ATXN3 in the same direction as in the primary screen.

**Figure 2.**
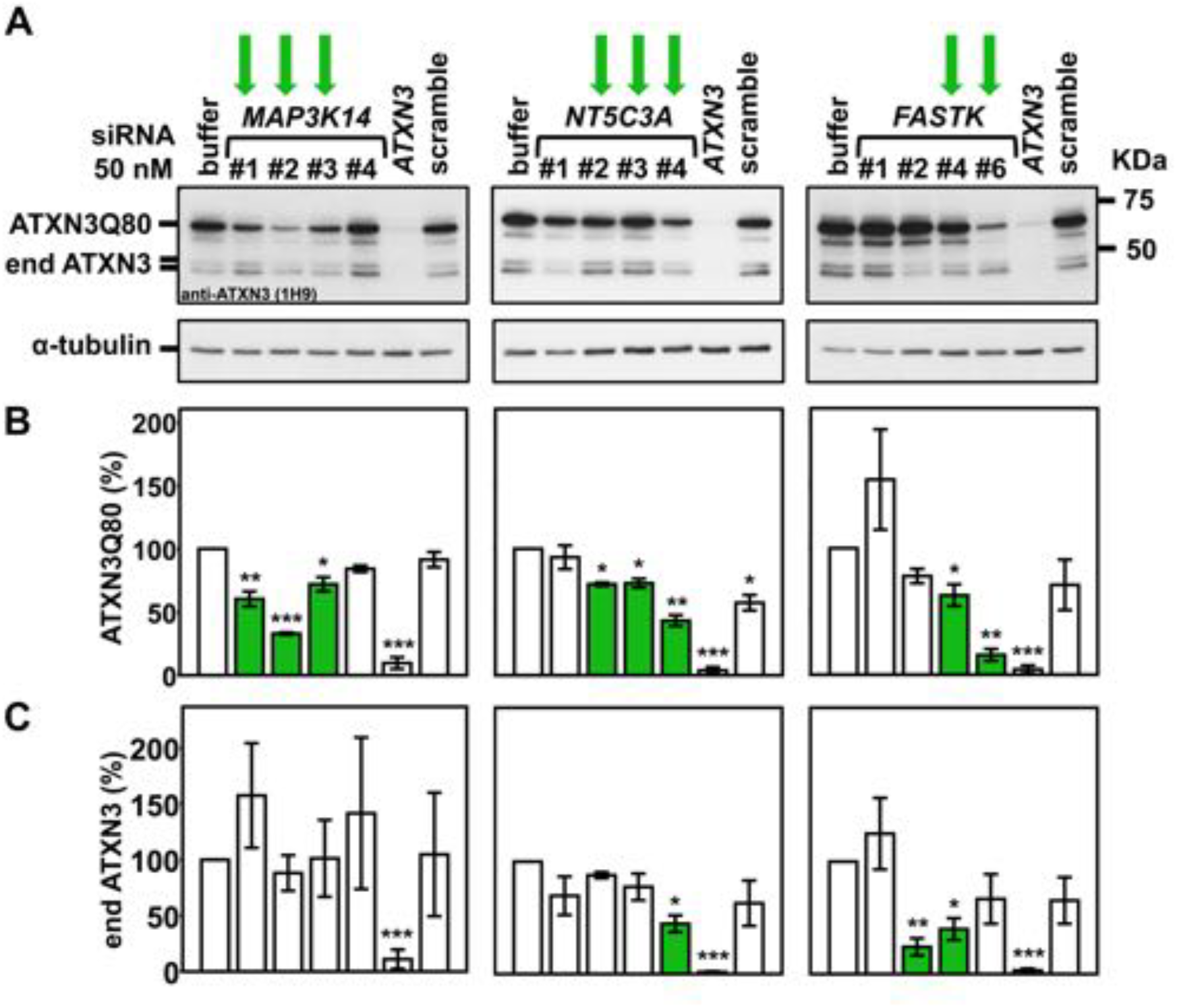
Three identified genes increase pathogenic ATXN3 levels in mammalian cells. A) Representative ATXN3 Western blots confirm the efficacy of two or more siRNAs targeting *MAP3K14*, *NT5C3*, and *FASTK* in decreasing levels of pathogenic FLAG-tagged human ATXN3Q80 in a stably expressing HEK293 cell line. Green arrows: siRNAs that effectively decrease ATXN3 levels. B, C) Histograms show quantification of ATXN3Q80 (B) and endogenous ATXN3 (end ATXN3) (C) from blots in (A) and another independent experiment. Bars represent the mean percentage of each protein relative to cells transfected with siRNA buffer alone, normalized for α-tubulin (± standard error of mean in two independent experiments). Green bars represent a statistically significant increase or decrease of ATXN3 levels compared to controls. * *P*<0.05, ** *P*<0.01, and *** *P*<0.001 are from student’s t-tests comparing each siRNA to siRNA buffer control.

**Figure 3.**
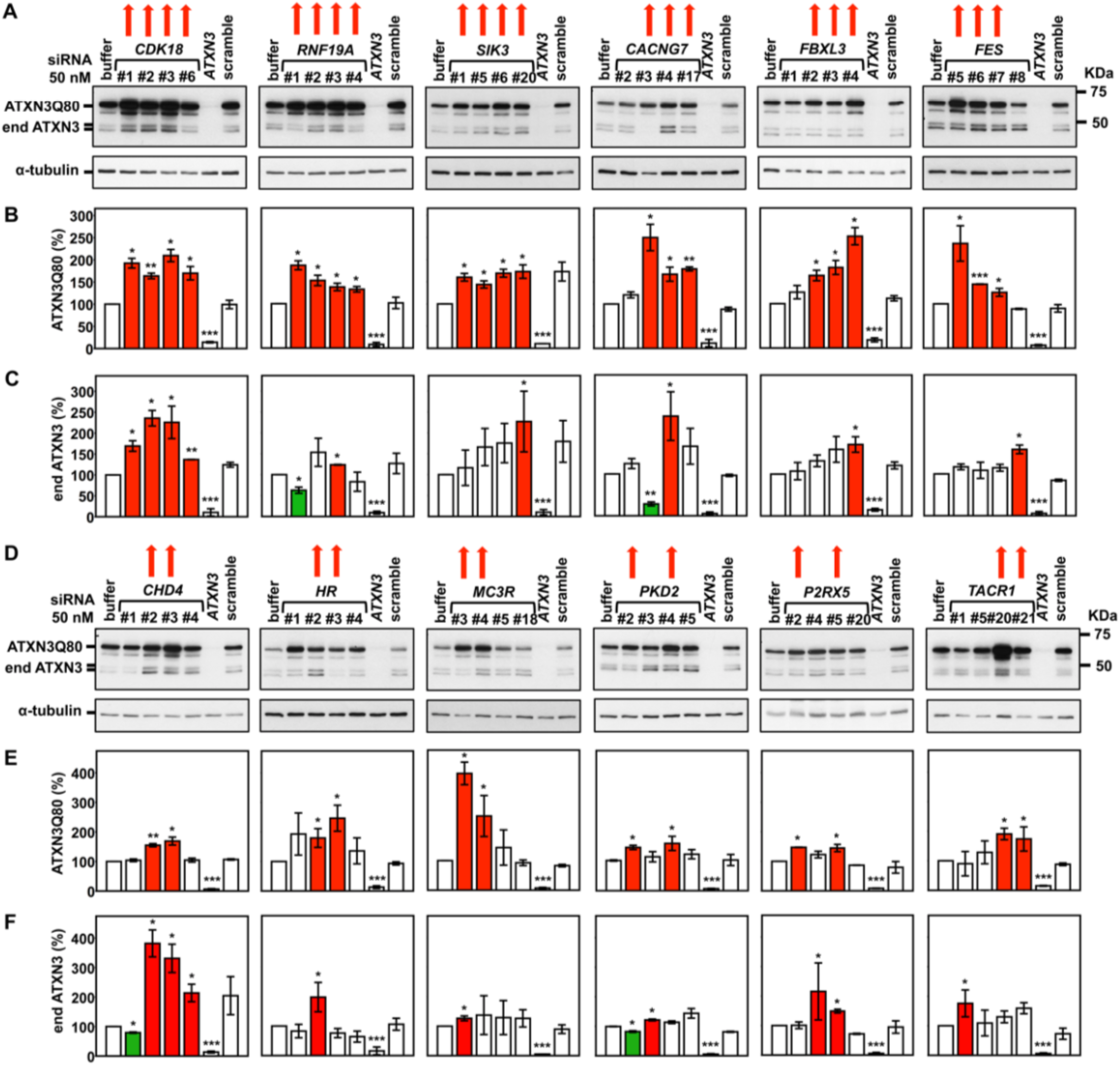
Twelve identified genes suppress pathogenic ATXN3 abundance in mammalian cells. A, D) Representative Western blots detecting ATXN3 reveal the efficacy of two or more siRNAs targeting *CDK8*, *RNF19A*, *SIK3*, *CACNG7*, *FBXL3*, *FES*, *CHD4*, *HR*, *MC3R*, *PKD2*, and *P2RX5*, *TACR1* in increasing levels of mutant ATXN3Q80. Red arrows: siRNAs that effectively increased ATXN3Q80 levels. B, C, E, F) Histograms showing the quantification of ATXN3Q80 (B, E) and endogenous ATXN3 (end ATXN3) (C, F) from blots in (A and D) and another independent experiment. Bars represent the mean percentage of each protein relative to cells transfected with siRNA buffer alone, normalized for α-tubulin (± standard error of mean in two independent experiments). Red and green bars represent, respectively, statistically significant increase or decrease of ATXN3 levels compared to controls. * *P*<0.05, ** *P*<0.01, and *** *P*<0.001 are from student’s t-tests comparing each siRNA to siRNA buffer control.

Using these criteria, we confirmed 15 of the 33 genes (Table 1): three enhancers of ATXN3 abundance (i.e. gene knockdown resulted in decreased ATXN3Q80 levels) – *MAP3K14*, *NT5C3A*, and *FASTK* (Figures 2A and B); and twelve suppressors of ATXN3 levels (i.e. gene knockdown resulted in increased ATXN3Q80 levels) – *CDK8*, *RNF19A*, *SIK3*, *CACNG7*, *FBXL3*, *FES*, *CHD4*, *HR*, *MC3R*, *PKD2*, *P2RX5*, and *TACR1* (Figures 3A, B, D and E). While most of these genes modulated the abundance of both expanded ATXN3Q80 and endogenous wild-type ATXN3, three preferentially regulated expanded ATXN3Q80 levels: *MAP3K14*, *RNF19A*, and *FES* (Figures 2 and 3). Bioinformatic analysis of these 15 genes revealed a potential molecular network with connections to tumor necrosis factor-α/nuclear factor-kappa B (TNF/NF-kB) and extracellular signal-regulated kinases 1 and 2 (ERK1/2) pathways (Figure 4).

**Figure 4.**
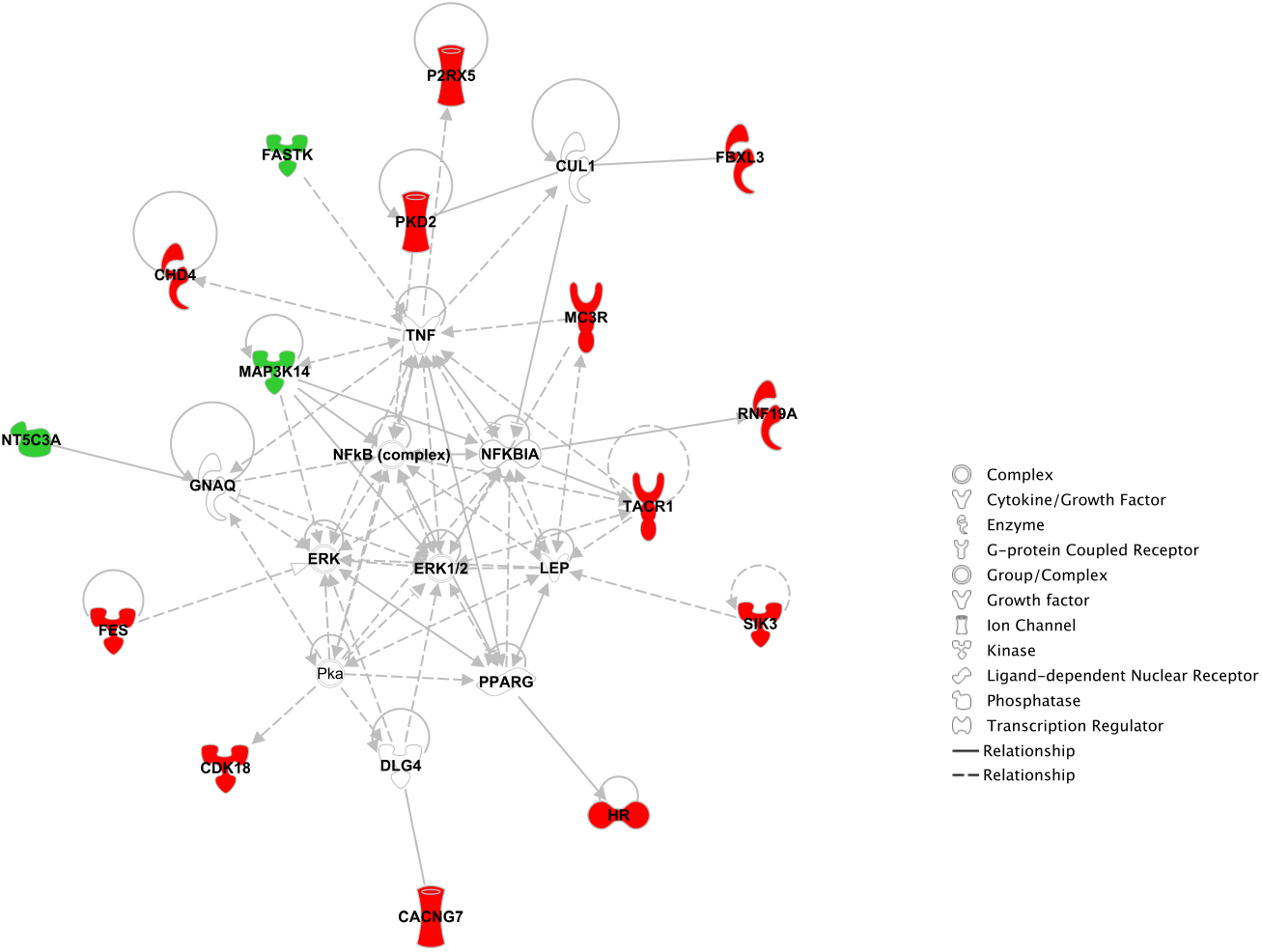
Molecular network formed by genes identified to modulate levels of pathogenic ATXN3 in HEK293 cells. IPA analysis of 15 genes reveals a molecular network with connections to TNF/NF-kB and ERK1/2 pathways. Genes whose knockdown decreased or increased ATXN3 levels are shown in green or red, respectively. Other genes relevant to the network but not identified as hits in our screen are don’t know depicted in grey. Legend for biological function of genes/proteins and gene relationships can be consulted at the left of network.

**Table 1.** Druggable genes that passed the secondary screen in ATXN3Q80 cells.

| **Gene Symbol** | **Entrez Gene Name** | **Entrez Gene ID** | **Cell location** | **Protein function** |
| --- | --- | --- | --- | --- |
| *CACNG7* | calcium voltage-gated channel auxiliary subunit gamma 7 | 59284 | Plasma Membrane | ion channel |
| *CDK18* | cyclin dependent kinase 18 | 5129 | Cytoplasm | kinase |
| *CHD4* | chromodomain helicase DNA binding protein 4 | 1108 | Nucleus | enzyme |
| *FASTK* | Fas activated serine/threonine kinase | 10922 | Cytoplasm | kinase |
| *FBXL3* | F-box and leucine rich repeat protein 3 | 26224 | Nucleus | enzyme |
| *FES* | FES proto-oncogene, tyrosine kinase | 2242 | Cytoplasm | kinase |
| *HR* | HR, lysine demethylase and nuclear receptor corepressor | 55806 | Nucleus | transcription regulator |
| *MAP3K14* | mitogen-activated protein kinase kinase kinase 14 | 9020 | Cytoplasm | kinase |
| *MC3R* | melanocortin 3 receptor | 4159 | Plasma Membrane | G-protein coupled receptor |
| *NT5C3A* | 5'-nucleotidase, cytosolic IIIA | 51251 | Cytoplasm | phosphatase |
| *P2RX5* | purinergic receptor P2X 5 | 5026 | Plasma Membrane | ion channel |
| *PKD2* | polycystin 2, transient receptor potential cation channel | 5311 | Plasma Membrane | ion channel |
| *RNF19A* | ring finger protein 19A, RBR E3 ubiquitin protein ligase | 25897 | Nucleus | enzyme |
| *SIK3* | SIK family kinase 3 | 23387 | Cytoplasm | kinase |
| *TACR1* | tachykinin receptor 1 | 6869 | Plasma Membrane | G-protein coupled receptor |

### Orthologs of *CHD4*, *FBXL3*, *HR* and *MC3R* regulate ATXN3Q77-induced toxicity in *Drosophila*

To assess whether the above findings in two cell models are physiologically relevant *in vivo*, we tested the efficacy of 10 orthologs of identified genes to alter mutant ATXN3-mediated toxicity in a *Drosophila* model of SCA3 [13, 28, 31]: *CACNG7*, *CHD4*, *CDK8*, *FASTK*, *FBXL3*, *FES*, *HR*, *MC3R*, *PKD2*, and *TACR1*. For simplicity, we focused on the fly eye expression model which is commonly used to examine the role and pathogenicity of various misfolded proteins [32]. Expression of pathogenic ATXN3 (Q77) in fly eyes is insufficiently toxic to cause marked degeneration of external structures [28, 31], necessitating the examination of internal eye structures for degenerative phenotypes or the use of a membrane-targeted GFP molecule (CD8-GFP) as a simple readout of the loss of the functional unit of the fly eye, the ommatidium [23]. Through this second assay, a toxic protein such as pathogenic ATXN3 is expressed in fly eyes independently of CD8-GFP; whereas the outside part of the fly eye seems unperturbed by the presence of the toxic protein, internal structures degenerate and photoreceptor cells disappear, resulting in the loss of GFP fluorescence [23]. Thus, increased GFP fluorescence reflects improved eye structure, whereas loss of fluorescence signifies internal eye structure degeneration [23, 25, 26]. In other words, this assay provides a quantifiable degenerative phenotype (reduced GFP) that facilitates screening [25]. In this model, UAS-CD8-GFP and UAS-ATXN3Q77 are driven independently by an eye-restricted driver, GMR-Gal4, through the binary Gal4-UAS system [33]. As shown in Figure 5A, expression of pathogenic ATXN3 in fly eyes leads to a statistically significant loss of GFP signal, as shown before [23].

**Figure 5.**
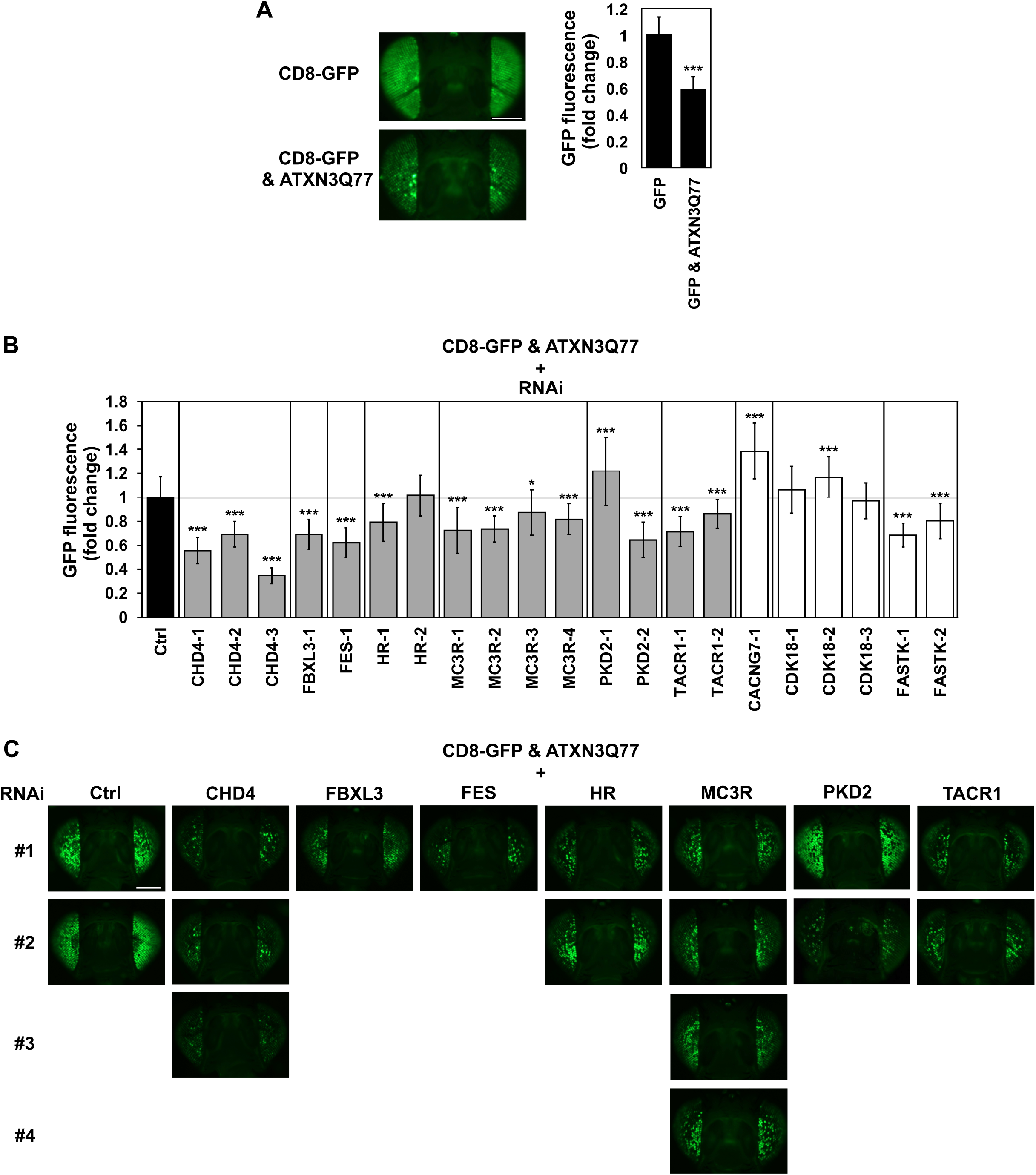
Effect of knockdown of specific fly genes on ATXN3-mediated degeneration. A) Expression of expanded ATXN3Q77 leads to degeneration in fly eyes, demonstrated by reduced CD8-GFP fluorescence. GMR-Gal4 was used to drive the independent expression of membrane-targeted GFP (CD8-GFP) and mutant ATXN3Q77 in fly eyes. Histograms on the right show quantification of GFP signal from images on the left and additional independent repeats. Scale bar: 200 µM. *** *P*<0.001 based on student’s t-test comparing GFP signal in the presence of ATXN3Q77 to signal in its absence. N ≥ 30 per genotype (note that these images were collected at a time and with a fluorescent bulb different than the ones in panel C). B) Quantification of the GFP signal from dissected fly heads that expressed pathogenic ATXN3Q77 as well as RNAi targeting the indicated genes. Numbers in RNAi lines indicate independent constructs. Grey bars highlight genes whose knockdown had an effect as expected, based on the observed modulation of ATXN3 levels in mammalian cells, whereas white bars highlight genes with opposite behavior compared to cell-based assays. Shown are means ± standard deviations. N ≥ 30 per genotype. * *P*<0.05, and *** *P*<0.001 are from student’s t-tests comparing each RNAi line to its respective control. C) Representative images of dissected fly heads expressing CD8-GFP alongside pathogenic ATXN3Q77 in the absence (Ctrl) or presence of RNAi targeting the noted genes (grey bars in (B)). Flies in all panels were seven days old. Numbers on the left side denote different RNAi transgenes used for targeted genes. All flies for the experiments shown here (C) and others that were used for quantification (B) were collected at the same time and imaged with the same fluorescent bulb. Scale bar: 200 µM.

For each of the 10 genes described above, we crossed RNAi *Drosophila* lines identified by BLAST analysis and FlyBase reports (Supplementary Table 3) to CD8-GFP & ATXN3Q77 flies, and then quantified GFP signal in dissected fly heads at day 7 (Figures 5B,C). RNAi targeting seven of 10 genes (orthologs of *CHD4*, *FBXL3*, *FES*, *HR*, *MC3R*, *PKD2*, and *TACR1*) resulted in statistically significant effects on ATXN3-mediated changes in GFP fluorescence in flt eyes (Figures 5B,C), consistent with the observed modulation of ATXN3 levels in mammalian cells (Figures 2 and 3), that is increased toxicity in fly eyes and increased levels of mutant ATXN3 in cells. In contrast, knockdown of orthologs of *CACNG7*, *CDK18* and *FASTK* in CD8-GFP & ATXN3Q77 flies led to results incongruent with cell-based data (Figure 5B). To evaluate the baseline effect of knockdown of these 10 genes on eye toxicity, we crossed the RNAi lines with CD8-GFP flies in the absence of ATXN3Q77 and observed the following compared with controls: 1) no differences on GFP signal for crosses with CHD4-1,2,3, FBXL3-1, MC3R-3, PKD2-1, TACR1-1,2 and FASTK-1 lines; 2) decreased GFP intensity in FES-1, HR-1,2, and MC3R-2,4 crosses; and 3) increased fluorescence in CACNG7-1, FASTK-2, CDK18-1,2,3 and PKD2-2 crosses (Supplementary Figure 5). Overall, the baseline toxicity of the RNAi lines on fly eyes did not interfere with the observed effect on ATXN3-mediated toxicity in crosses of CD8-GFP & ATXN3Q77 flies.

We next evaluated the effect of these seven genes on ATXN3-mediated disruption of internal eye structures (Figure 6A). Histological analyses of eye sections confirmed that knockdown of the orthologs for four genes (*CHD4*, *FBXL3*, *HR*, and *MC3R)* enhanced ATXN3Q77 toxicity (Figure 6A), highlighted by increased separation of retinal structures from the underlying lamina. While no apparent differences in eye structure were observed in crosses of ATXN3Q77 with RNAi lines for orthologs of *FES*, *PKD2* and *TACR1*, fly heads from all seven crosses showed increased levels of ATXN3Q77 protein (Figure 6B and Supplementary Figure 6), in accordance with our expectations from the cell-based assays described above (Figure 3). Among the seven orthologs for which we confirmed modulation of ATXN3Q77-mediated toxicity, *CHD4*, *FBXL3*, *HR*, and *MC3R* surfaced as the top toxicity suppressor genes to pursue further because gene knockdown resulted in concordant outcomes in all three readouts of toxicity or protein abundance in flies (fluorescence intensity, histology and Western blot).

**Figure 6.**
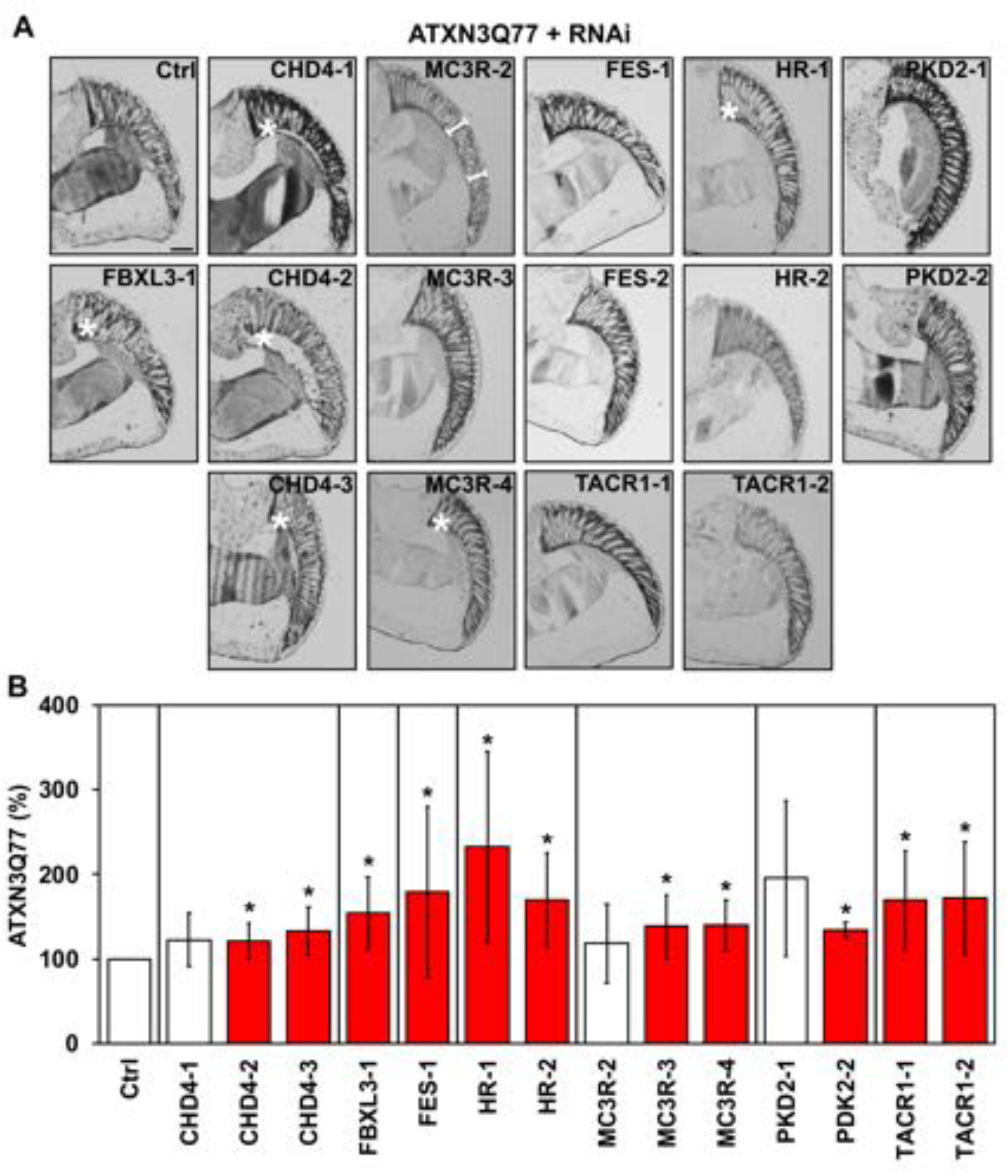
Depletion of four fly genes increases ATXN3-dependent toxicity in fly eyes. A) Representative images of histological sections from fly eyes expressing pathogenic ATXN3Q77 in the absence (Ctrl) or presence of RNAi constructs targeting the indicated genes. Numbers on the left indicate that more than one RNAi line was used for each gene. Scale bar: 50 µM. White asterisks (*) highlight separation of basal retinal structures. White brackets highlight shortening of ommatidial length. B) Graph showing the quantification of ATXN3Q77 protein Western blot bands detected by anti-MJD antibody (Supplementary Figure 6) relative to controls in fly heads from crosses of ATXN3Q77 flies with RNAi lines for selected genes, normalized to total protein levels measured by Direct Blue 71. Bars show means ± standard deviations. N ≥ 3 independent experimental repeats. Red histograms show increased levels of ATXN3 compared to controls. * *P*<0.05 is from student’s t-tests comparing each RNAi line to its respective control.

### Overexpression of *FBXL3* suppresses ATXN3 abundance in a CUL1-dependent manner in SCA3 neuronal progenitor cells (NPCs)

We selected *FBXL3* to further confirm its role in regulating mutant ATXN3 abundance in human cells expressing pathogenic ATXN3 from the endogenous locus, namely SCA3 hESC-derived NPCs. *FBXL3* was chosen for further analysis because its protein is directly implicated in mechanisms of protein degradation. *FBXL3* encodes a F-box protein that is a component of the ubiquitin protein ligase complex SKP1-Cullin1-F-box (SCF) involved in ubiquitin-dependent protein degradation [34]. We first generated NPCs from control and SCA3 hESCs [35] and confirmed that these cells express the markers of neural progenitor lineage PAX6, SOX1 and Nestin (Figure 7A). In confirming ATXN3 expression in these cells by immunofluorescence, we observed increased ATXN3-positive puncta in the nucleus and cytoplasm of SCA3 NPCs compared to control NPCs (Figure 7B).

**Figure 7.**
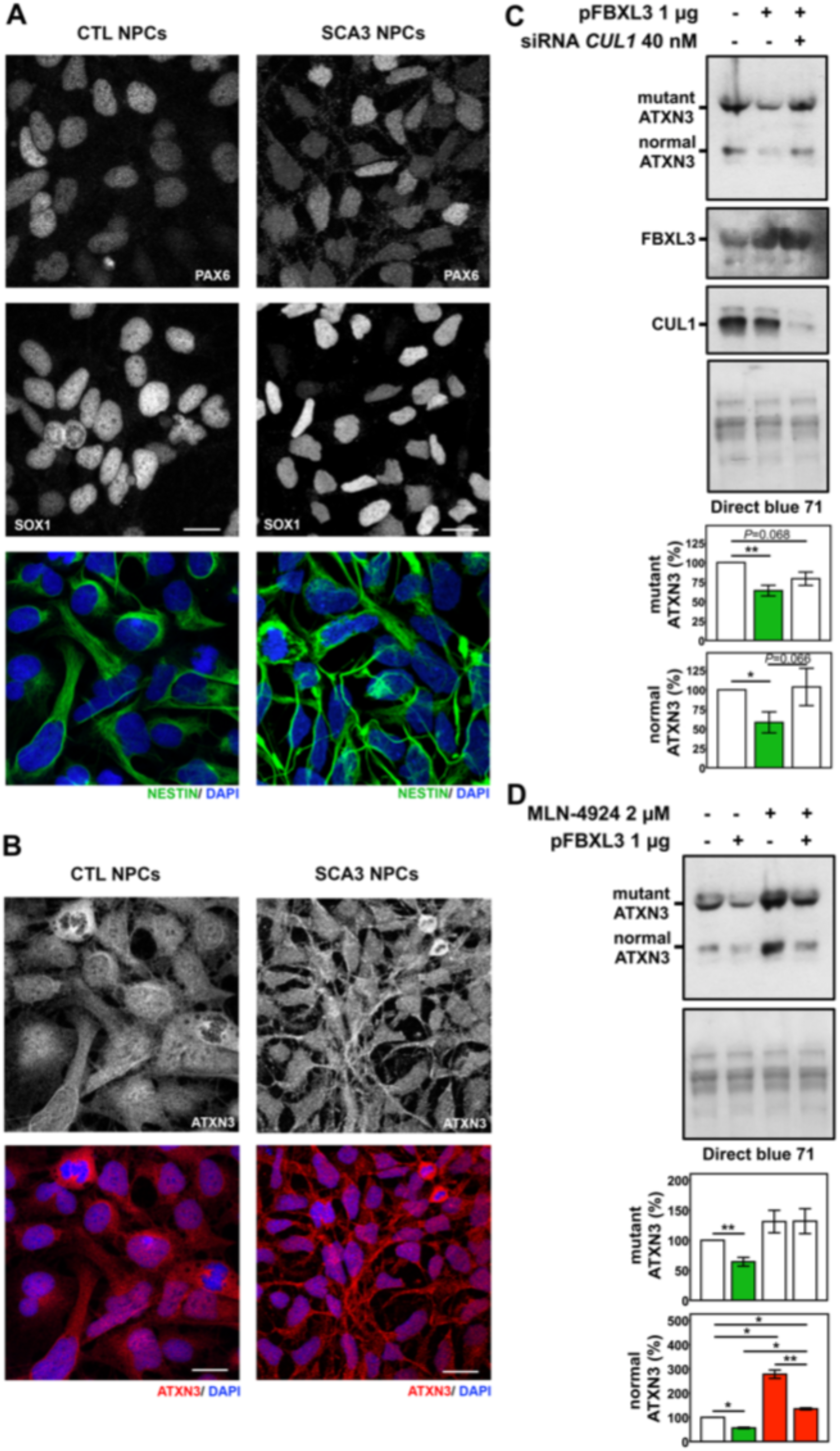
FBXL3 suppresses ATXN3 levels in SCA3 neuronal progenitor cells (NPCs). A) Control (CTRL) and SCA3 NPCs showing immunostaining for NPC markers PAX6 (white), SOX1 (white) and Nestin (green). B) CTRL and SCA3 NPCs immunostained for ATXN3 (red) using anti-MJD antibody. Nuclei were stained with DAPI (blue). Photographs are 2 μm z-stacks acquired by confocal imaging. Scale bar: 25 μM. C) Western blots detecting ATXN3 (anti-MJD) in protein extracts of SCA3 NPCs overexpressing FBXL3, with or without concomitant siRNA-mediated knockdown of CUL1 for 48 hours. D) Representative immunoblot blot detecting ATXN3 (anti-MJD) in SCA3 NPCs overexpressing FBXL3 for 48 hr and treated with 2 μM of CUL1 inhibitor MLN-4924 for the final 16 hours. Quantification of bands corresponding to mutant and normal ATXN3 are shown in the accompanying graphs. Bars represent the mean percentage of ATXN3 relative to mock-electroporated cells and normalized to total protein levels measured by Direct Blue 71 (± SEM) in three independent experiments. Red and green bars represent, a statistically significant increase or decrease, respectively, of ATXN3 levels compared to controls. * *P*<0.05 is from one-tailed *t*-test comparing the different conditions.

We overexpressed FBXL3 in SCA3 NPCs and confirmed that high levels of FBXL3 reduce endogenous levels of both wild-type and mutant ATXN3 proteins to 58% and 64%, respectively, of control levels (Figure 7C). To evaluate if FBXL3-mediated reduction of ATXN3 abundance occurs via the SCF/CUL1 ubiquitination complex, we co-electroporated plasmid overexpressing FBXL3 while also decreasing CUL1 with siRNAs against CUL1 (Figure 7C). Knockdown of CUL1 abolished FBXL3-mediated reduction of normal ATXN3, but only accounted for about half of the observed FBXL3-facilitated decrease of pathogenic ATXN3 (Figure 7C), suggesting that FBXL3 handles or recognizes normal and mutant ATXN3 differently. Overexpression of FBXL3 and knockdown of CUL1 were confirmed at the transcript level (Supplementary Figure 7). ATXN3 transcript levels, although variable across experiments, were actually higher when FBXL3 was overexpressed, with or without knockdown of CUL1 (Supplementary Figure 7) indicating that the observed FBXL3-mediated reduction of ATXN3 levels occurs at the protein rather than transcriptional level, as expected.

To further explore the role of FBXL3/SCF in modulating ATXN3 protein levels we treated SCA3 NPCs with MLN-4924, an inhibitor of Cullin-RING E3 ubiquitin ligase (CRL) activation, in the presence or absence of FBXL3 overexpression. Baseline MLN-4924 treatment increased levels of wild-type ATXN3 to 277% of control levels and showed a trend, albeit not statistically significant, to increase mutant ATXN3 to 131% of controls (Figure 7D). This result implies that SCF and CRL complexes mediate normal ATXN3 clearance, presumably via ubiquitin-dependent degradation, to a greater extent than mutant ATXN3 clearance. In addition, the effect of MLN-4924 on normal ATXN3 was largely countered by overexpressing FBXL3 (134% of control levels) (Figure 7D). Collectively, while these results implicate SCF and CRL complexes as regulators of ATXN3 protein levels; they also suggest that the action of FBXL3 on ATXN3 may be multifaceted. The fact that FBXL3 can counter some of the effects of inhibiting SCF and CRL on wild-type ATXN3, but not pathogenic ATXN3, raises the possibilty of differential regulation of the two forms of ATXN3.

## Discussion

There are currently no disease-modifying therapies for SCA3. Reducing levels of mutant *ATXN3* transcript or encoded protein, however, has effectively mitigated disease phenotypes in preclinical trials in SCA3 transgenic mouse models [10–19, 36]. Accordingly, therapeutic approaches that deplete pathogenic ATXN3 proteins in the SCA3 brain appear promising. While some evidence suggests that expanded polyQ ATXN3 can be degraded by the proteasome [37–42] or macroautophagy [43, 44] and that its stability is affected by specific protein interactions [38, 45, 46], we lack comprehensive knowledge of the pathways controlling the abundance of mutant ATXN3. Because ATXN3 is a deubiquitinating enzyme that participates in ubiquitin-dependent protein quality control pathways [47–50], the way cells handle this particular protein could be unusually complex. This knowledge prompted the unbiased druggable genome siRNA screen reported here, which identified several genes as regulators of ATXN3 protein abundance. Because proteins encoded by druggable genes can be inhibited or activated by drugs, the genes discovered here represent compelling therapeutic targets in SCA3 and possibly other polyQ diseases. Our additional studies of one identified gene, *FBXL3*, also offer new insights into the cellular pathways by which ATXN3 is likely degraded.

Employing an iterative screening platform that leveraged a broad range of methods to detect changes in pathogenic ATXN3 levels and toxicity, we successively identified: i) 33 of 2742 druggable genes as specific modulators of ATXN3 using a cell-based ATXN3-Luc assay; ii) 15 of 33 genes whose knockdown significantly decreased or increased levels of pathogenic ATXN3 protein in a secondary SCA3 cell model; iii) seven of 10 ortholog genes in *Drosophila* whose knockdown increased mutant ATXN3-mediated toxicity, assessed by fluorescence signal in eyes of CD8-GFP & ATXN3Q77 flies; and iv) four of seven fly ortholog genes whose knockdown increased mutant ATXN3 abundance and showed disrupted internal eye structures in ATXN3Q77 flies. Of these four genes we then selected one, *FBXL3*, for further mechanistic studies, which showed that *FBXL3* regulates wild-type and pathogenic ATXN3 levels in human SCA3 NPCs, primarily via a SCF complex-dependent pathway.

Our primary screen identified 317 genes equally distributed through the categories of enhancers (N=163) and suppressors (N=154) of ATXN3-Luc signal in controls. These genes distributed similarly over different protein function categories, implying that a variety of classes of proteins are involved at some level with handling ATXN3. This is not an unexpected outcome since ATXN3 has been implicated in various cellular processes through its DUB activity [3, 47–52]. Among the genes that arose from secondary assays, fifteen are related to TNF/NF-kB and ERK1/2 pathways, indicating that pathogenic ATXN3 levels may be affected by TNF- or mitogen-dependent signaling. TNF, a cytokine mainly produced by glial cells in the brain, either promotes inflammation through the NF-kB pathway and apoptotic cell death, or is neuroprotective, depending on the precise receptors it binds to [53]. The mitogen-activated protein kinases (MAPKs) ERK1/2, also connected with the TNF/NF-kB pathway, likewise can either promote neuronal survival or neuronal death [54]. Glia is understudied in SCA3, but several recent findings suggest key roles for glia and inflammatory signaling in SCA3: early transcriptional changes in SCA3 mouse oligodendrocytes [55], the contribution of astrocyte-like glia to non-cell autonomous degeneration in SCA3 flies [56], and neuroprotection from the NSAID ibuprofen in a SCA3 mouse model [57]. Our findings that pathogenic ATXN3 protein levels can be regulated by TNF/NF-kB and ERK1/2 pro-inflammatory and cell death/survival pathways highlight the need for further investigation of their role in SCA3.

Genes and proteins that regulate pathogenic ATXN3 abundance and toxicity yet have a limited number of substrates and/or effectors would be ideal targets for intervention in SCA3. Among such candidates, the F-box protein FBXL3, which binds substrates and promotes their ubiquitination and subsequent degradation [34], peeked our interest. Our observation that FBXL3 regulates levels of endogenous wild-type and pathogenic ATXN3 in human SCA3 NPCs, supports the view that ATXN3 is a substrate for FBXL3 under physiological conditions reflective of the human disease. While only a few FBXL3 substrates have been validated, 141 proteins were recently identified as being potentially recruited to SCF^FBXL3^ complexes by cryptochromes CRY1 and CRY2, which are themselves FBXL3 substrates [58–60]. It remains, however, to be determined whether targeting FBXL3 will prove to be a viable therapeutic strategy in SCA3. On the one hand, null *FBXL3* mutations cause autosomal recessive developmental delay and intellectual disability [61] suggesting that FBXL3 mediates the stability of numerous proteins and that complete loss of FBXL3 function is harmful for cells, at least during development. On the other hand, *FBXL3* knockdown in our mammalian cell lines and in flies did not show accompanying toxicity (Supplementary Table 2 and Supplementary Figure 5). Further studies will be needed to establish whether increasing or decreasing FBXL3 has effects on the fully developed adult brain, which is the relevant target in age-related neurodegenerative diseases such as SCA3.

If *FBXL3* knockdown increases ATXN3 abundance, then its overexpression would be expected to decrease ATXN3 protein levels. Exogenous expression of FBXL3 in SCA3 NPCs indeed reduced the levels of both wild-type and pathogenic ATXN3. The potential role of the SCF and potentially other CRL complexes in ubiquitinating and regulating ATXN3 levels appears to span to both wild-type and pathogenic forms of this DUB. Supporting evidence for the particular involvement of the SCF complex in ATXN3 ubiquitination comes from a recent report showing that CUL1 and FBXO33 specifically interact and promote ubiquitination and solubility of a truncated form of pathogenic ATXN3 [62], and from an independent yeast-two-hybrid study that identified the mouse homologs Cul1 and Atxn3 as interacting partners (Costa et al. unpublished observations). As ATXN3 shows both DUB and deneddylase activities *in vitro* [63], future work should investigate its functions in pathways regulated by SCF and CRL complexes and the effect that these complexes have on ATXN3.

## Conclusions

We identified 15 druggable genes that implicate the involvement of TNF- or mitogen-dependent signaling cascades in regulating pathogenic ATXN3 levels. One of these potential candidates for SCA3 intervention, *FBXL3*, is directly involved in protein quality control and was effective in modulating the levels of mutant and wild-type ATXN3 under physiological conditions in human cells. The proteins encoded by these genes and the pathways in which they are implicated demand further evaluation to understand the pathobiology of SCA3 and to seek disease-modifying therapies for this fatal disorder.

## List of Abbreviations

ATXN3: ataxin-3
CRL: Cullin-RING E3 ubiquitin ligase
DUB: deubiquitinating enzyme
GFP: green fluorescent protein
hESC: human embryonic stem cell
MJD: Machado-Joseph disease
NPC: neuronal progenitor cell
PBS: phosphate buffered saline
polyQ: polyglutamine
RIPA: radioimmunoprecipitation assay buffer
RT: room temperature
SCA3: Spinocerebellar ataxia type 3
SCF: SKP1-Cullin1-F-box
SEM: standard error of the mean

## Declarations

### Ethics approval

All human embryonic stem cells studies were approved by the Human Pluripotent Stem Cell Research Oversight (HPSCRO) of the University of Michigan (application 1097).

### Consent for publication

Not applicable.

### Availability of data and materials

All data generated or analyzed during this study are included in this article and its supplementary information files.

### Authors’ Contributions

MCC, HLP and SVT designed the study. NSA, JRS, YY, BR, KL, EDS, AJB, SVT and MCC executed experiments. NSA, JRS, SVT and MCC analyzed data. MCC prepared the figures and drafted the manuscript. SVT and HLP reviewed the manuscript critically.

### Authors’ Information

NSA, MSc, Research Lab Technician Lead, currently applying to a Physician Assistant Program

JRS, Ph.D., Ph.D. student, currently Pharmaceutical Industry Scientist

YY, BSc, Research Assistant, currently Pharm. D. Resident

BR, BSc, Undergraduate Researcher, currently applying to medical schools

KL, M.D., Research Associate

EDS, BSc, Research Assistant

AJB, BSc, Research Laboratory Technician Associate

SVT, Ph.D., Associate Professor & Associate Dean

HLP, M.D., Ph.D., Professor

MCC, Ph.D., Research Assistant Professor

## Acknowledgements

The authors thank Steve Swanson and Martha J. Larsen from the Center of Chemical Genomics at the University of Michigan for assistance with high-throughput siRNA screen, Dr. Richard McEachin for bioinformatics support, and Dr. Garry Smith and Laura Keller for initial training with hESCs and NPCs.

