## Supplementary Tables and Figures for "Druggable genome screen identifies new regulators of the abundance and toxicity of ATXN3, the Spinocerebellar Ataxia Type 3 disease protein"

**Supplementary Table 1: Primers used to assess transcript levels by quantitative RT-PCR.**

**Supplementary Table 2: Summary of luminescence inhibition and viability for the 33 genes whose knockdown was selective for ATXN3-Luc assay.**

**Supplementary Table 3: Fly orthologue RNAi lines.**

**Supplementary Figure 1: Map of assay plates in the primary druggable genome screen.** Legend is located at the bottom of the plate schematic.

**Supplementary Figure 2: Distribution of the 317 hits from primary screen by protein function and subcellular localization.** Number of hits per class is displayed in each slice/division of the pie chart. Legend of pie charts is located at the right of the respective chart.

**Supplementary Figure 3: Confirmation of gene knockdown of the three identified enhancers of mutant ATXN3 levels.** Histograms represent transcript levels for *MAP3K14*, *NT5C3*, or *FASTK* siRNA transfections of ATXN3Q80 HEK293 cells parallel to experiments depicted in Figure 2. Transcript fold change for each gene was normalized for *ACTB* expression and referenced to the average of siRNA buffer-treated cells. Bars represent the average of transcript fold change per siRNA treatment (± SEM in two independent experiments). * $P<0.05$, ** $P<0.01$, and *** $P<0.001$ are from student's t-tests comparing each siRNA to siRNA buffer control. Green arrows: siRNAs that effectively decrease pathogenic ATXN3 protein levels (Figure 2).

**Supplementary Figure 4: Confirmation of gene knockdown of the twelve identified suppressors of mutant ATXN3 levels.** Histograms represent transcript levels for *CDK8*,

*RNF19A*, *SIK3*, *CACNG7*, *FBXL3*, *FES*, *CHD4*, *HR*, *MC3R*, *PKD2*, *P2RX5*, or *TACR1* siRNA transfections of ATXN3Q80 HEK293 cells parallel to experiments depicted in Figure 3. Transcript fold change for each gene was normalized for *ACTB* expression and referenced to the average of siRNA buffer-treated cells. Bars represent the average of transcript fold change per siRNA treatment (± SEM in two independent experiments). * $P$<0.05, ** $P$<0.01, and *** $P$<0.001 are from student's t-tests comparing each siRNA to siRNA buffer control. Red arrows: siRNAs that effectively increase mutant ATXN3 protein levels (Figure 3).

**Supplementary Figure 5: Effect of knockdown of specific fly genes on CD8-GFP flies.** Quantification of the GFP signal from dissected fly heads that expressed RNAi targeting the indicated genes. Numbers in RNAi lines indicate independent constructs. As in Figure 5, grey bars highlight genes whose gene knockdown in CD8-GFP & ATXN3Q77 had an effect as expected, based on the observed modulation of ATXN3 levels in mammalian cells, whereas white bars highlight genes with opposite behavior compared to cell-based assays. Shown are means ± standard deviations. N ≥ 13 per genotype. ** $P$<0.01, and *** $P$<0.001 are from student's t-tests comparing each RNAi line to its respective control.

**Supplementary Figure 6: Representative Western blot detecting ATXN3 in fly heads from crosses of ATXN3Q77 flies with RNAi lines for selected genes.** Quantification of ATXN3Q77 protein Western blot bands detected by anti-MJD antibody is shown in Figure 5B relative to controls and normalized to total protein levels measured by Direct Blue 71. N ≥ 10 fly heads per lane.

**Supplementary Figure 7: Transcript levels of *FBXL3*, *CUL1* and *ATXN3* in SCA3 NPCs overexpressing *FBXL3*, with or without simultaneous *CUL1* knockdown.** Histograms represent transcript levels for each gene in each transfection condition of SCA3 NPCs performed in parallel to experiments depicted in Figure 7. Transcript fold change for each gene was normalized for *ACTB* expression and referenced to the average of mock-treated cells. Bars represent the average of transcript fold change per transfection condition (± SEM in three independent experiments). * $P<0.05$ is from student's t-tests comparing each transfection condition to mock control.

**Supplementary Figure 1**

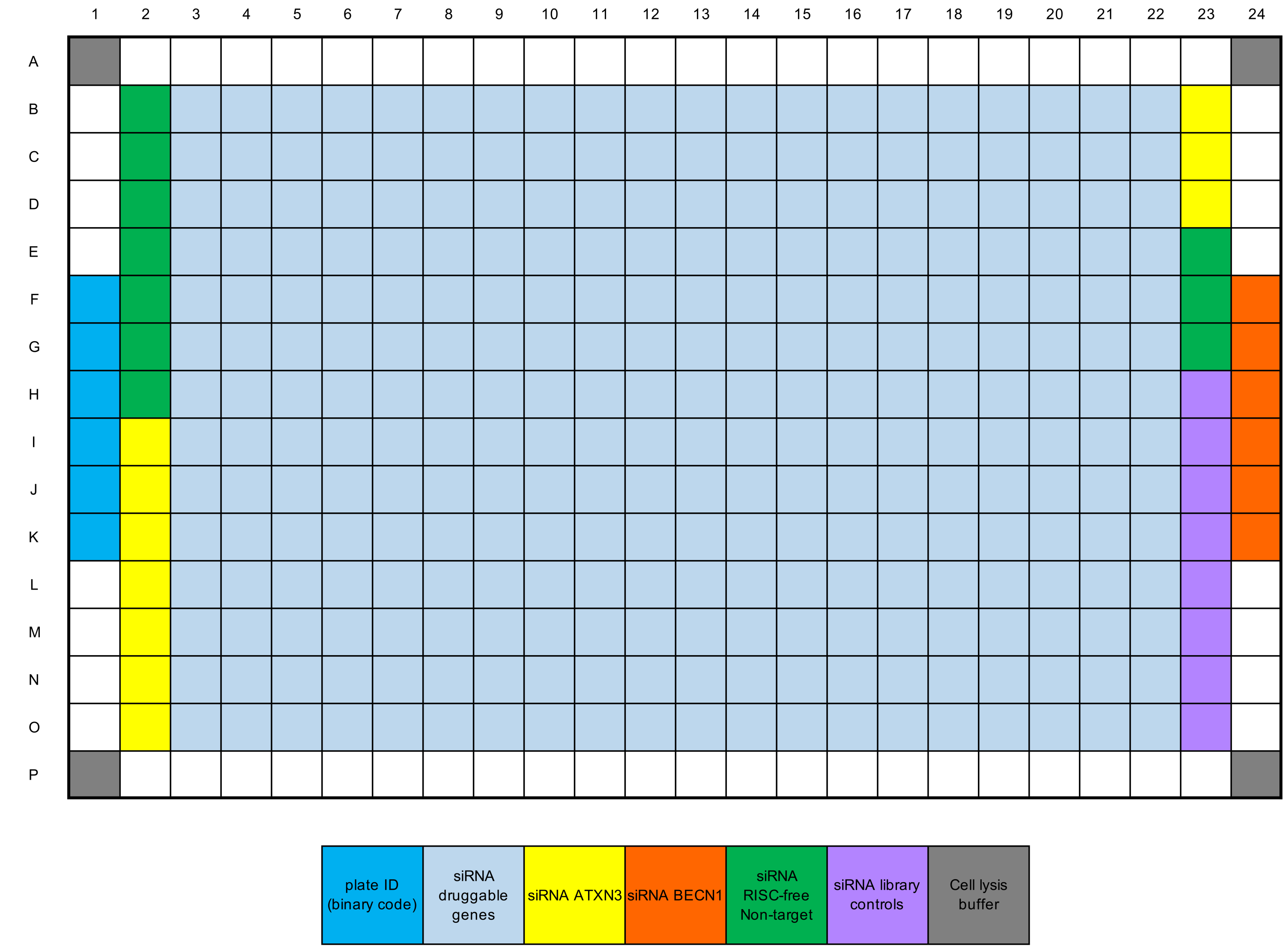

**Supplementary Figure 2**

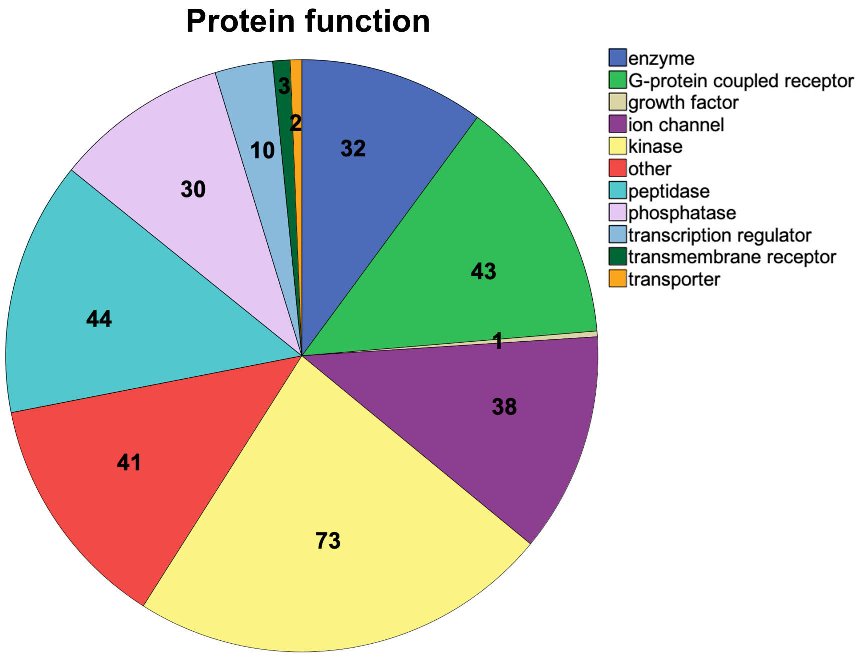


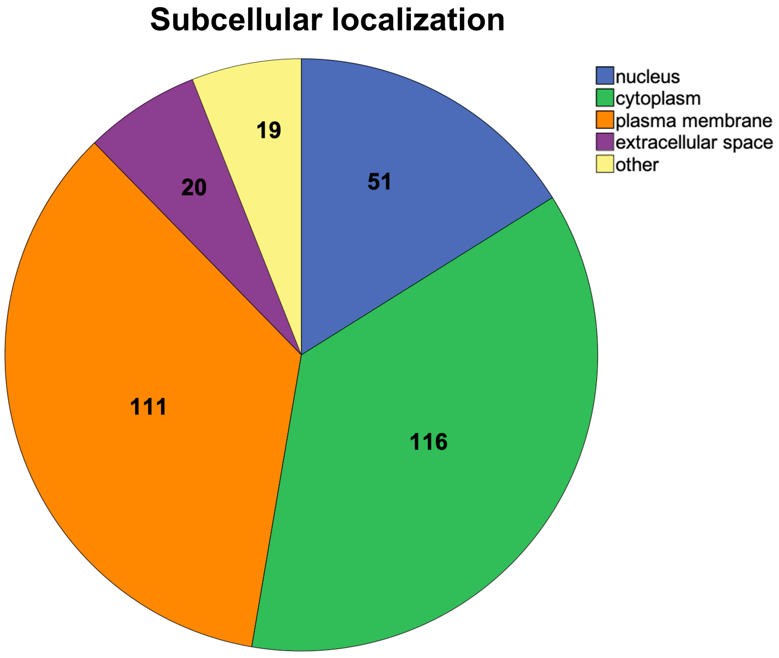

**Supplementary Figure 3**

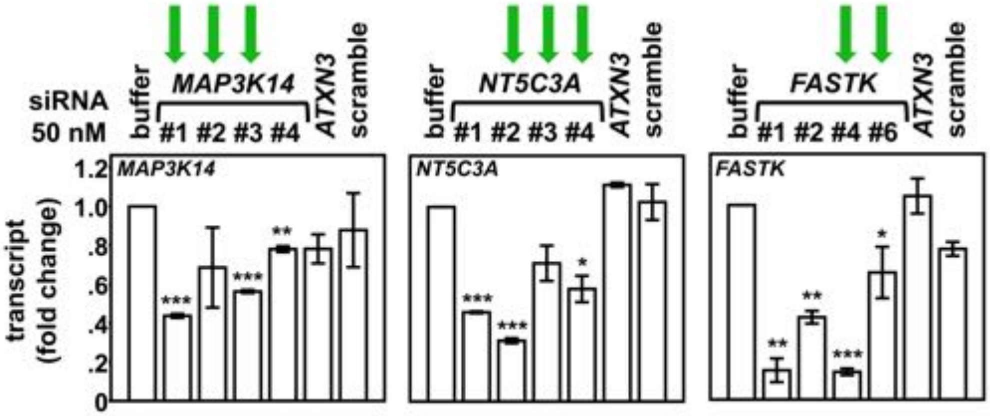

Supplementary Figure 4

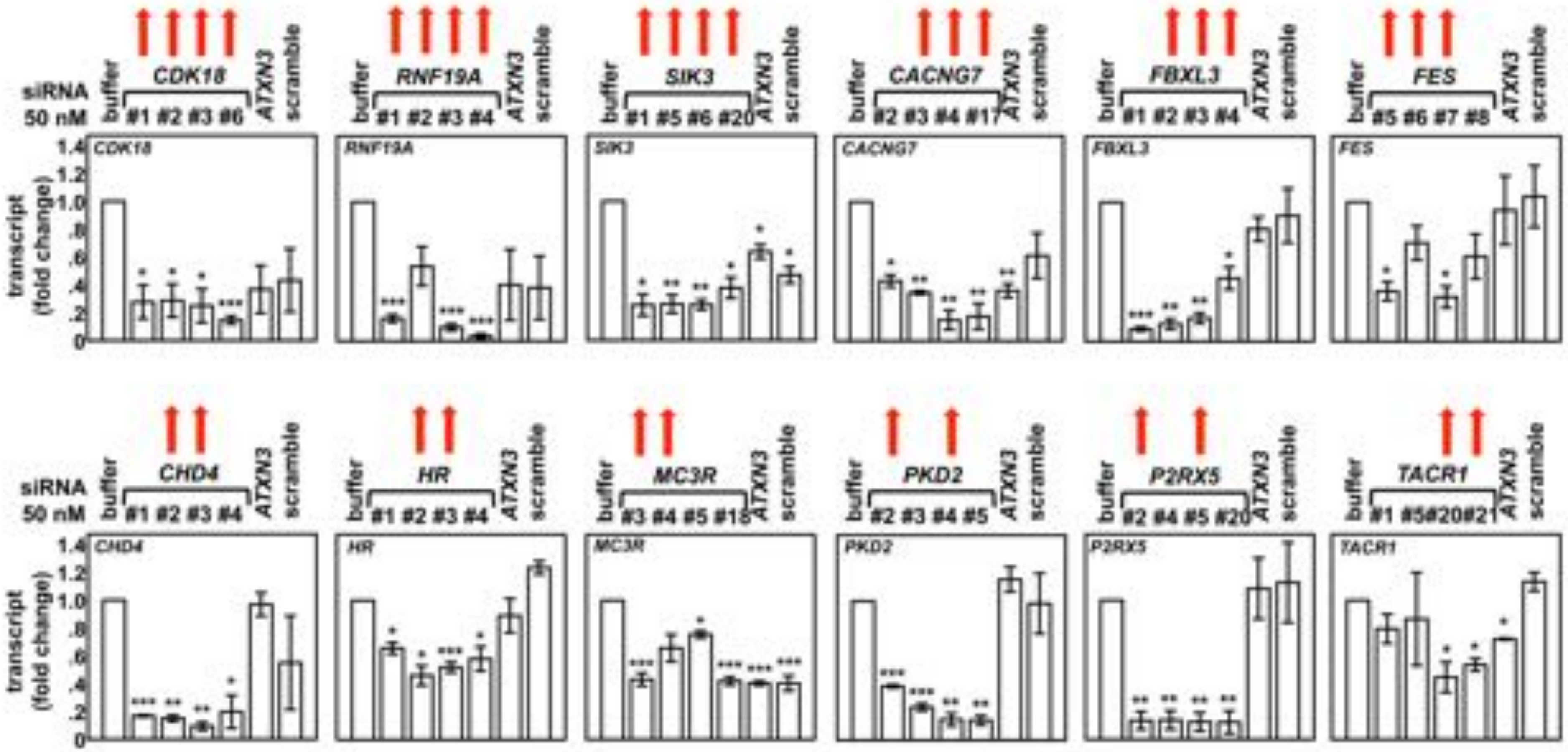

#### Supplementary Figure 5

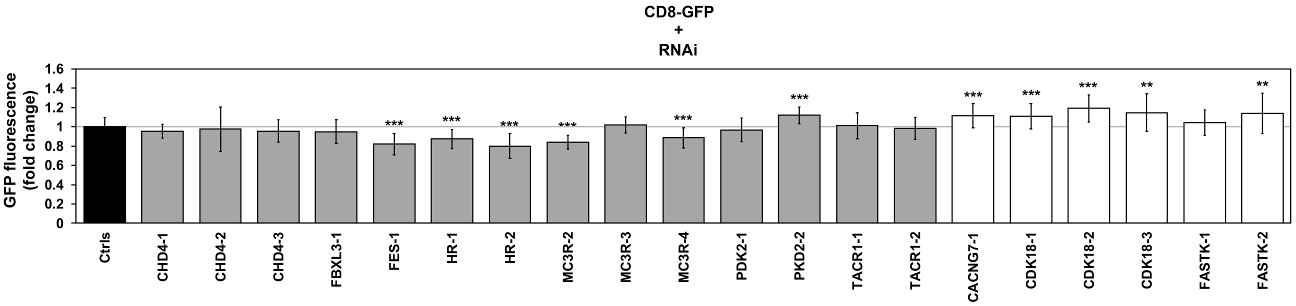

**Supplementary Figure 6**

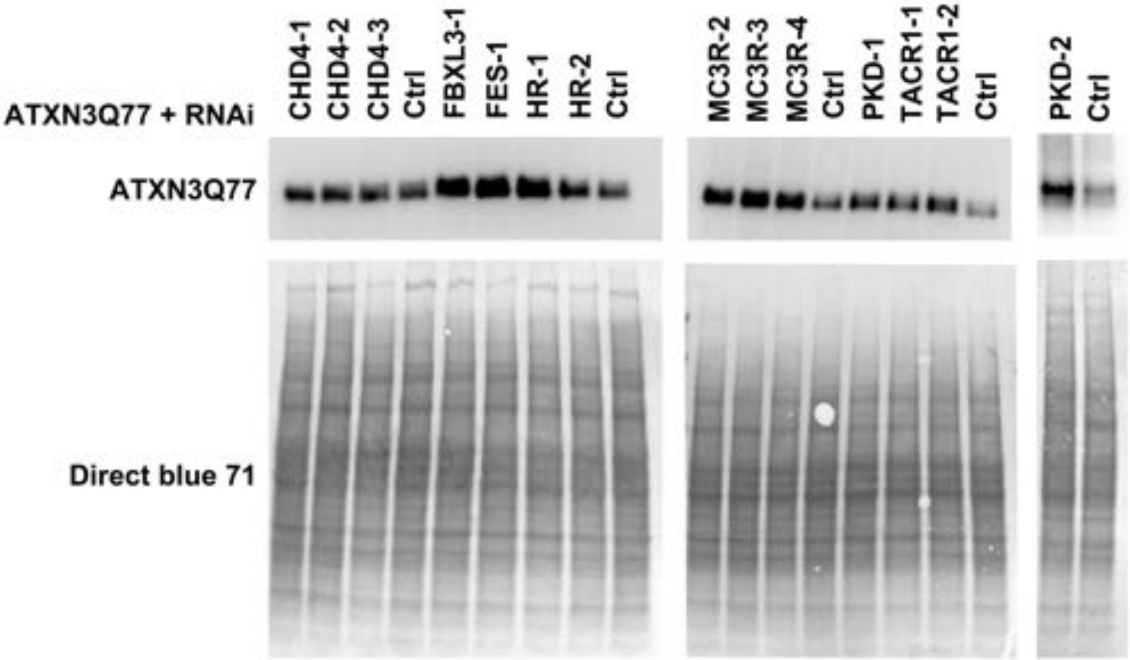

### Supplementary Figure 7

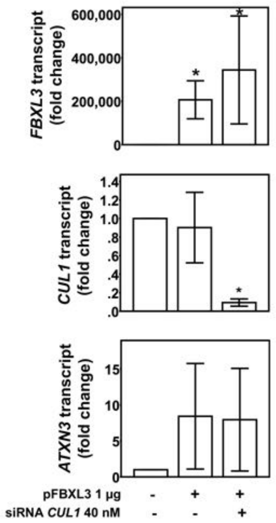

**Supplementary Table 1**. Primers used to assess transcript levels by quantitative RT-PCR.

| Gene | Primer | |
|---|---|---|
| | Name | Sequence 5' -> 3' |
| *ATCB* | ATCB_F1 | CGTCCACACCCGCCG |
| | ATCB_F2 | CCACCATCACGCCCTGG |
| *ATXN3* | hATXN3_F3 | CGAGAAACAAGAAGGCTCACTT |
| | hATXN3_R3 | CCCAAACTTTCAAGGCATTGCT |
| *CACNG7* | CACNG7_F3 | TTCTTTGCAGGTCGGGAGAAA |
| | CACNG7_R3 | ACGAGGAAGAGGCTGACCAT |
| *CDK18* | CDK18_F1 | CGCCGGGTATAAGGAGCAAA |
| | CDK18_R1 | CTGCAAGTTCTCATTCCGCC |
| *CHD4* | CHD4_F1 | CCACCGAATCCTCAACCACA |
| | CHD4_R1 | CACCCCTCATTAACTCCCTGT |
| *CUL1* | CUL1_F1 | CAGGGTCTTGCAGCCATTGA |
| | CUL1_R1 | AAGCGACCACAAGCCTTATCA |
| *FASTK* | FASTK_F1 | AAGTGCCCATCTTTCCCCAG |
| | FASTK_R1 | CAGCAGGAAGTCTGTGCAGT |
| *FBXL3* | FBXL3_F1 | AGCTTCAAGGTGGACAGCAG |
| | FBXL3_R1 | ACGAACACAACTGTCAGTGC |
| *FES* | FES_F1 | CTGGTGTTGGGTGAGCAGAT |
| | FES_R1 | TGCTTCAGGATCCTCGCTTC |
| *HR* | HR_F1 | CCAGCCGTGACAGAGGAC |
| | HR_R1: 5’ | GGACTGCTCCTGAAAGCCTG |
| *KCNA7* | KCNA7_F1 | GCGCCGCGAGTATTTCTTCG |
| | KCNA7_R1 | CGTAGAAGGCCACCTCTTCCA |
| *MC3R* | MC3R_F1 | GCGACTACCTGACCTTCGAG |
| | MC3R_R1 | TAGCGGAGCGCGTAAAAGAT |
| *MAP3K14* | MAP3K14_F2 | AGTACAGCCAGTCCGAGAGT |
| | MAP3K14_R2 | GCCAGGGACTTTGAGCTCTT |
| *NT5C3A* | NT5C3A_F2 | TGCTTGGACGCGCATGA |
| | NT5C3A_R2 | ATGATCTTGGTCTTCCGCCC |
| *P2RX5* | P2RX5_F2 | TGGAAACGGAGTGAAGACCG |
| | P2RX5_R2 | GGGGAAACGGATGTGGTTCT |
| *PKD2* | PKD2_F1 | TTGTGCATCTTGACCTACGGC |
| | PKD2_R1 | AGCCCATCCAATAAGGAGCC |
| *RNF19A* | RNF19A_F1 | GGAGTCTGTCAGGAAGTGCC |
| | RNF19A_R1 | GCCTTCTTTGTCCAATGGGATG |
| *TACR1* | TACR1_F1 | CCCATCATCTACTGCTGCCT |
| | TACR1_R1 | CCAGGCGGCTGACTTTGTA |

**Supplementary Table 2.** Summary of luminescence inhibition and viability for the 33 genes whose knockdown was selective for ATXN3-Luc assay.

| Gene Symbol | GENEID | Gene Accession | GINumber | Dharmacon siRNA pool catalog Number | Dharmacon individual siRNA catalog number | Primary screen ATXN3-Luc cells - siRNA smart pool for each gene | | | Confirmation screen ATXN3-Luc cells - 4 individual siRNAs for each gene | |
|---|---|---|---|---|---|---|---|---|---|---|
| | | | | | | activity | luminescence inhibition (%) | cell viability (%) | luminescence inhibition (%) | cell viability (%) |
| *C2* | 717 | NM_000063 | 20631970 | MU-005797-01 | D-005797-01 | increase luminescence levels | -131.73 | 105.21 | -129.90 | 96.90 |
| | | | | | D-005797-03 | | | | -39.90 | 80.15 |
| | | | | | D-005797-04 | | | | -24.63 | 103.31 |
| | | | | | D-005797-17 | | | | -34.33 | 105.08 |
| *CHD4* | 1108 | NM_001273 | 51599155 | MU-009774-01 | D-009774-01 | increase luminescence levels | -104.63 | 85.72 | -80.67 | 96.42 |
| | | | | | D-009774-02 | | | | -159.13 | 119.28 |
| | | | | | D-009774-03 | | | | -354.40 | 68.98 |
| | | | | | D-009774-04 | | | | | |
| *CHRM5* | 1133 | NM_012125 | 38176159 | MU-005466-00 | D-005466-01 | decrease luminescence levels | 62.67 | 86.91 | | |
| | | | | | D-005466-02 | | | | | |
| | | | | | D-005466-03 | | | | 62.25 | 60.21 |
| | | | | | D-005466-04 | | | | 35.55 | 61.70 |
| *DAPK3* | 1613 | NM_001348 | 4557510 | MU-004947-00 | D-004947-01 | decrease luminescence levels | 62.43 | 91.49 | | |
| | | | | | D-004947-02 | | | | | |
| | | | | | D-004947-03 | | | | | |
| | | | | | D-004947-04 | | | | 49.85 | 61.72 |
| *FES* | 2242 | NM_002005 | 13376997 | MU-003130-01 | D-003130-05 | increase luminescence levels | -129.33 | 93.49 | | |
| | | | | | D-003130-06 | | | | -93.07 | 101.06 |
| | | | | | D-003130-07 | | | | -93.33 | 74.31 |
| | | | | | D-003130-08 | | | | | |
| *KCNA7* | 3743 | NM_031886 | 25952091 | MU-006217-00 | D-006217-01 | decrease luminescence levels | 70.43 | 98.22 | 39.75 | 66.97 |
| | | | | | D-006217-02 | | | | | |
| | | | | | D-006217-03 | | | | | |
| | | | | | D-006217-04 | | | | 33.95 | 95.16 |
| *MC3R* | 4159 | NM_019888 | 17986278 | MU-005659-02 | D-005659-03 | increase luminescence levels | -117.73 | 96.53 | | |
| | | | | | D-005659-04 | | | | -245.13 | 112.91 |
| | | | | | D-005659-05 | | | | -59.60 | 101.69 |
| | | | | | D-005659-18 | | | | | |
| *MST1* | 4485 | NM_020998 | 31543211 | MU-008946-00 | D-008946-01 | decrease luminescence levels | 59.03 | 95.93 | | |
| | | | | | D-008946-02 | | | | | |
| | | | | | D-008946-03 | | | | 46.40 | 97.45 |
| | | | | | D-008946-04 | | | | | |
| *NTRK1* | 4914 | NM_001007792 | 56118209 | MU-003159-02 | D-003159-05 | decrease luminescence levels | 60.10 | 90.24 | | |
| | | | | | D-003159-06 | | | | 34.53 | 121.54 |
| | | | | | D-003159-07 | | | | | |
| | | | | | D-003159-08 | | | | | |
| *P2RX5* | 5026 | NM_175081 | 28416936 | MU-006286-04 | D-006286-02 | increase luminescence levels | -111.97 | 95.94 | -90.17 | 84.03 |
| | | | | | D-006286-04 | | | | -325.30 | 121.83 |
| | | | | | D-006286-05 | | | | | |
| | | | | | D-006286-20 | | | | -176.80 | 113.83 |
| *CDK18* | 5129 | NM_002596 | 38505200 | MU-004836-02 | D-004836-01 | increase luminescence levels | -128.67 | 87.15 | -308.67 | 86.67 |
| | | | | | D-004836-02 | | | | -186.53 | 104.22 |
| | | | | | D-004836-03 | | | | -101.67 | 92.25 |
| | | | | | D-004836-06 | | | | -78.37 | 99.28 |
| *PFKFB4* | 5210 | NM_004567 | 19923257 | MU-006764-01 | D-006764-01 | decrease luminescence levels | 73.50 | 95.54 | | |
| | | | | | D-006764-02 | | | | | |
| | | | | | D-006764-04 | | | | | |
| | | | | | D-006764-17 | | | | 40.45 | 117.98 |
| *PGK1* | 5230 | NM_000291 | 22095338 | MU-006767-01 | D-006767-01 | decrease luminescence levels | 74.70 | 102.36 | | |
| | | | | | D-006767-02 | | | | | |
| | | | | | D-006767-04 | | | | 80.95 | 100.77 |
| | | | | | D-006767-05 | | | | | |
| *PKD2* | 5311 | NM_000297 | 33286447 | MU-006288-01 | D-006288-02 | increase luminescence levels | -149.53 | 100.48 | -223.50 | 93.51 |
| | | | | | D-006288-03 | | | | -56.07 | 97.00 |
| | | | | | D-006288-04 | | | | -240.17 | 82.81 |
| | | | | | D-006288-05 | | | | -22.40 | 102.75 |
| *PTPN13* | 5783 | NM_006264 | 5453991 | MU-008065-00 | D-008065-01 | increase luminescence levels | -102.13 | 97.78 | | |
| | | | | | D-008065-02 | | | | -113.57 | 78.36 |
| | | | | | D-008065-03 | | | | -25.97 | 102.81 |
| | | | | | D-008065-04 | | | | -550.27 | 108.30 |
| *TACR1* | 6869 | NM_015727 | 7669545 | MU-005733-03 | D-005733-01 | increase luminescence levels | -107.07 | 89.18 | | |
| | | | | | D-005733-05 | | | | -41.57 | 117.00 |
| | | | | | D-005733-20 | | | | -186.40 | 89.35 |
| | | | | | D-005733-21 | | | | -35.83 | 106.08 |
| *MAP3K14* | 9020 | NM_003954 | 115298644 | MU-003580-04 | D-003580-10 | decrease luminescence levels | 72.87 | 106.23 | | |
| | | | | | D-003580-11 | | | | 73.80 | 80.56 |
| | | | | | D-003580-12 | | | | | |
| | | | | | D-003580-26 | | | | | |
| *FASTK* | 10922 | NM_006712 | 39995105 | MU-005317-02 | D-005317-01 | decrease luminescence levels | 64.57 | 98.71 | | |
| | | | | | D-005317-02 | | | | | |
| | | | | | D-005317-04 | | | | | |
| | | | | | D-005317-06 | | | | 76.05 | 71.12 |
| *IRAK3* | 11213 | NM_007199 | 6005791 | MU-004762-00 | D-004762-01 | decrease luminescence levels | 64.70 | 91.63 | | |
| | | | | | D-004762-02 | | | | | |
| | | | | | D-004762-03 | | | | | |
| | | | | | D-004762-04 | | | | 41.35 | 87.57 |
| *TWF2* | 11344 | NM_007284 | 40068460 | MU-003169-01 | D-003169-05 | decrease luminescence levels | 80.67 | 99.78 | 63.25 | 96.34 |
| | | | | | D-003169-06 | | | | | |
| | | | | | D-003169-07 | | | | | |
| | | | | | D-003169-08 | | | | | |
| *KDM2A* | 22992 | NM_012308 | 16306579 | MU-012458-00 | D-012458-01 | increase luminescence levels | -127.20 | 98.40 | -224.40 | 107.91 |
| | | | | | D-012458-02 | | | | -291.23 | 101.11 |
| | | | | | D-012458-03 | | | | -368.00 | 100.69 |
| | | | | | D-012458-04 | | | | -47.37 | 107.34 |
| *DCUN1D4* | 23142 | NM_015115 | 94536779 | MU-014118-02 | D-014118-01 | increase luminescence levels | -144.17 | 102.31 | | |
| | | | | | D-014118-02 | | | | -75.47 | 99.91 |
| | | | | | D-014118-03 | | | | -232.90 | 101.37 |
| | | | | | D-014118-04 | | | | -201.73 | 115.82 |
| *SIK3* | 23387 | NM_025164 | 39812204 | MU-004779-03 | D-004779-01 | increase luminescence levels | -113.10 | 102.75 | -213.40 | 109.78 |
| | | | | | D-004779-05 | | | | -69.10 | 105.42 |
| | | | | | D-004779-06 | | | | -257.23 | 109.27 |
| | | | | | D-004779-20 | | | | -98.33 | 119.34 |
| *RNF19A* | 25897 | NM_015435 | 35493781 | MU-006965-00 | D-006965-01 | increase luminescence levels | -175.20 | 97.11 | -352.20 | 109.35 |
| | | | | | D-006965-02 | | | | -67.53 | 95.56 |
| | | | | | D-006965-03 | | | | -46.67 | 95.32 |
| | | | | | D-006965-04 | | | | -37.97 | 104.07 |
| *FBXL3* | 26224 | NM_012158 | 16306583 | MU-012422-00 | D-012422-01 | increase luminescence levels | -123.47 | 94.47 | | |
| | | | | | D-012422-02 | | | | -141.77 | 99.47 |
| | | | | | D-012422-03 | | | | -84.17 | 112.51 |
| | | | | | D-012422-04 | | | | -240.37 | 108.48 |
| *PRICKLE4* | 29964 | NM_013397 | 118722346 | MU-006979-01 | D-006979-01 | increase luminescence levels | -110.33 | 107.36 | | |
| | | | | | D-006979-02 | | | | | |
| | | | | | D-006979-03 | | | | -85.77 | 93.44 |
| | | | | | D-006979-04 | | | | -310.10 | 97.47 |
| *EMR2* | 30817 | NM_152918 | 23397686 | MU-005488-02 | D-005488-01 | decrease luminescence levels | 59.80 | 96.56 | 28.50 | 139.39 |
| | | | | | D-005488-02 | | | | | |
| | | | | | D-005488-19 | | | | | |
| | | | | | D-005488-20 | | | | | |
| *NT5C3* | 51251 | NM_016489 | 70608210 | MU-032020-01 | D-032020-01 | decrease luminescence levels | 62.40 | 95.06 | | |
| | | | | | D-032020-02 | | | | | |
| | | | | | D-032020-03 | | | | | |
| | | | | | D-032020-04 | | | | 55.30 | 62.67 |
| *HR* | 55806 | NM_018411 | 70906481 | MU-011872-01 | D-011872-01 | increase luminescence levels | -111.87 | 95.08 | -142.33 | 95.00 |
| | | | | | D-011872-02 | | | | -95.50 | 104.44 |
| | | | | | D-011872-03 | | | | -99.53 | 92.68 |
| | | | | | D-011872-04 | | | | -143.73 | 114.13 |
| *SMURF1* | 57154 | NM_181349 | 31317289 | MU-007191-01 | D-007191-01 | decrease luminescence levels | 64.20 | 95.58 | | |
| | | | | | D-007191-02 | | | | 48.95 | 68.42 |
| | | | | | D-007191-03 | | | | | |
| | | | | | D-007191-21 | | | | | |
| *CACNG7* | 59284 | NM_031896 | 22027498 | MU-017347-01 | D-017347-02 | increase luminescence levels | -110.40 | 99.16 | -29.37 | 96.86 |
| | | | | | D-017347-03 | | | | -195.23 | 97.57 |
| | | | | | D-017347-04 | | | | -116.37 | 105.40 |
| | | | | | D-017347-17 | | | | -28.47 | 97.81 |
| *FBXL20* | 84961 | NM_032875 | 14249619 | MU-015029-00 | D-015029-01 | decrease luminescence levels | 69.97 | 94.18 | 44.85 | 88.27 |
| | | | | | D-015029-02 | | | | | |
| | | | | | D-015029-03 | | | | | |
| | | | | | D-015029-04 | | | | | |
| *PHF13* | 148479 | NM_153812 | 24432092 | MU-016306-00 | D-016306-01 | increase luminescence levels | -102.27 | 92.17 | | |
| | | | | | D-016306-02 | | | | | |
| | | | | | D-016306-03 | | | | -87.27 | 127.59 |
| | | | | | D-016306-04 | | | | -51.13 | 148.00 |

**Supplementary Table 3.** Fly orthologue RNAi lines.

| Human Gene | Fly Orthologue | RNAi line Stock Number | Provenance |
|---|---|---|---|
| *CDK18* | CG10579 (Eip63E) | 35579, 28901, 34075 | Bloomington *Drosophila* Stock Center |
| *CHD4* | CG8103 (Mi-2) | 35398, 51774, 33419 | Bloomington *Drosophila* Stock Center |
| *HR* | CG8165 (JHDM2) | 32975, 58264 | Bloomington *Drosophila* Stock Center |
| *TACR1* | CG7887 (TkR99D) | 27513, 55732 | Bloomington *Drosophila* Stock Center |
| *FASTK* | CG31643 (no assigned *Drosophila* symbol) | 53380, 56938 | Bloomington *Drosophila* Stock Center |
| *KCNA7* | CG12348 (Sh) | 31680, 53347 | Bloomington *Drosophila* Stock Center |
| *FES* | CG8874 (FER) | 33375 | Bloomington *Drosophila* Stock Center |
| *FBXL3* | CG9952 (Ppa) | 31357 | Bloomington *Drosophila* Stock Center |
| *PKD2* | CG8808 (Pdk) | 28635, 35142 | Bloomington *Drosophila* Stock Center |
| *CACNG7* | CG33670 (stg1) | 41928 | Bloomington *Drosophila* Stock Center |
| *MC3R* | *CG18741 (Dop1R2)* CG9652 (Dop1R1) | *26018, 51423,* 55239, 31765 | Bloomington *Drosophila* Stock Center |

Order of RNAi stocks matches the RNAi (#) in figures.

For MC3R, two fly genes matched; we chose RNAi lines for both.

Italics on Fly Orthologs indicate which RNAi lines correspond to which gene for MC3R.
